# Quantitative fate mapping: Reconstructing progenitor field dynamics via retrospective lineage barcoding

**DOI:** 10.1101/2022.02.13.480215

**Authors:** Weixiang Fang, Claire M. Bell, Abel Sapirstein, Soichiro Asami, Kathleen Leeper, Donald J. Zack, Hongkai Ji, Reza Kalhor

## Abstract

Natural and induced somatic mutations that accumulate in the genome during development record the phylogenetic relationships of cells; however, whether these lineage barcodes can capture the dynamics of complex progenitor fields remains unclear. Here, we introduce quantitative fate mapping, an approach to simultaneously map the fate and quantify the commitment time, commitment bias, and population size of multiple progenitor groups during development based on a time-scaled phylogeny of their descendants. To reconstruct time-scaled phylogenies from lineage barcodes, we introduce Phylotime, a scalable maximum likelihood clustering approach based on a generalizable barcoding mutagenesis model. We validate these approaches using realistically-simulated barcoding results as well as experimental results from a barcoding stem cell line. We further establish criteria for the minimum number of cells that must be analyzed for robust quantitative fate mapping. Overall, this work demonstrates how lineage barcodes, natural or synthetic, can be used to obtain quantitative fate maps, thus enabling analysis of progenitor dynamics long after embryonic development in any organism.

## Introduction

Embryonic development is the genesis of complex body plans in the animal kingdom. It starts with the zygote, a single cell in a totipotent state, and ends with millions of specialized terminal cells organized in tissues. In between, dividing cells assume increasingly diverse but decreasingly potent intermediate progenitor states. Each progenitor state specifies the ensuing states that its cells may take thus directing them toward their terminal fates. Collectively, progenitor states orchestrate the formation of complex tissues by ensuring the emergence of all terminal cell types in harmony. Therefore, delineating how progenitor states relate to each other and terminal fates—the cell fate map—is critical for our understanding of normal and dysregulated development as well as our ability to generate engineered tissues. However, mapping cell fate is challenging due to the number of progenitor states involved, their interdependent relationships, and the time lapse between terminal fates and most of the progenitor states that help create them.

The recent advances in genome engineering and sequencing have inspired a new approach for interrogating cell fates during development: retrospective lineage analysis using synthetic or natural somatic barcodes [1,2]. These approaches rely on the somatic accumulation of random mutations in the genome during development. Each mutation is inherited by the descendants of the cell in which it occurs; each descendant can add new mutations to the combination it inherited. This process marks each terminal cell with a combination of mutations—a barcode—that encodes its phylogenetic relationship to the other cells [3]. Synthetic lineage barcoding, which relies on gene editing technologies to induce mutations, has been implemented in a variety of model systems including the zebrafish [4–7], fruit fly [8], and mouse [9–11]. Natural lineage barcoding, which relies on naturally-accumulating somatic mutations, has been primarily used in humans [12–15]. And while the mechanism, timing, and extent of mutagenesis differ between various implementations, the functional outcome remains the same: genomic barcodes that can retrospectively inform cell phylogeny within one organism at a single-cell resolution. As cells’ fate decisions are also somatically inherited to their daughters through epigenetic mechanisms [16], these retrospective approaches hold unique promise for mapping cell fate because, unlike single-cell molecular profiling approaches, they can bridge time lapses between terminal cells and their progenitors that exist far earlier. Moreover, unlike prospective lineage tracing approaches, they allow parallel analysis of multiple progenitor groups and do not depend on the identification and manipulation of progenitors.

Despite this compelling potential, the full scope of the information that retrospective lineage barcoding can provide about the fate of the intermediate progenitor states remains unclear for several reasons. First, cell phylogeny is a function of cell divisions and most cell divisions in higher organisms do not accompany fate decisions. In the roundworm *C. elegans*, a unique model system in which almost all cell divisions give rise to daughters with different fates, the phylogeny of terminal cells is identical to the fate of their progenitors [17]. In more complex organisms, on the other hand, cell fate is often determined at the level of progenitor populations which also undergo cell divisions that are not associated with fate decisions, leading to divergences between phylogeny and fate [2,18,19] (**Figure 1**). As a result, adjacent liver hepatocytes of identical progenitor state history may have the maximum possible distance on the phylogenetic tree by being the descendants of different cells at the 2-cell stage. Second, while the progenitor states and their fate remain largely stereotyped within species, the phylogenetic histories of the cell populations that assume those progenitor states can vary greatly from embryo to embryo due to stochasticity in fate decisions [14]. Third, single-cell lineage barcodes can be obtained for only a fraction of the cell population due to the practical limitations of single-cell sequencing. Whereas most mammals have millions of cells in each tissue, current technologies can only sequence thousands of single cells. Given the divergences between fate and phylogeny and the variable nature of the latter, it remains unclear how phylogenies derived from small samples can reliably inform organism-level fate maps. These considerations raise critical questions about the value of measuring cell phylogeny through barcoding approaches in complex organisms: What features of progenitor populations in embryonic development are reflected in the phylogeny of sampled terminal cells? Can retrospective barcodes, synthetic or natural, be used to extract these features and if so how?

**Figure 1.**
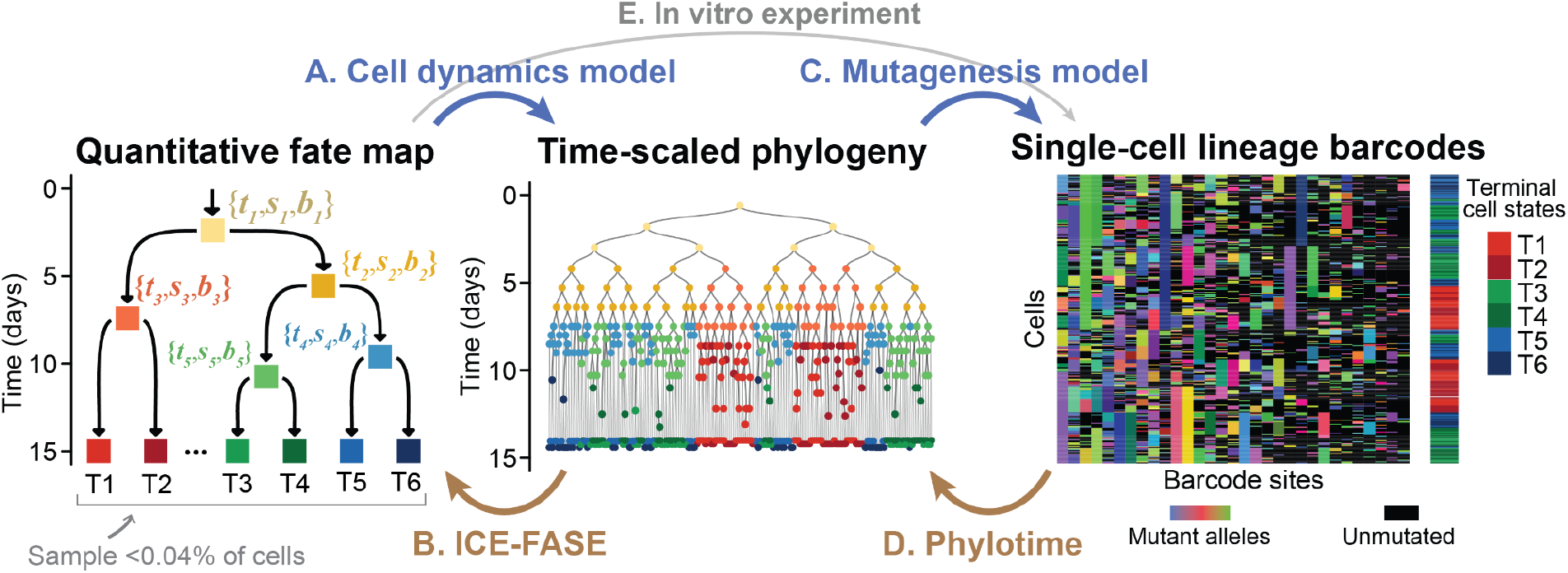
Graphical outline of this study, establishing quantitative fate mapping. Quantitative fate maps specify progenitor state dynamics and are used to generate time-scaled cell phylogenies and lineage barcoding results in terminal cells (blue arrows). Phylotime and ICE-FASE enable quantitative fate mapping by reconstructing first time-scaled phylogenies and then quantitative fate maps from lineage barcode data (brown arrows). *{t*_*i*_, *s*_*i*_, *b*_*i*_*}* are dynamic parameters of each progenitor state: commitment time, population size, and commitment bias. An in vitro system is used to experimentally verify model assumptions (gray arrow and brown arrows).

To address these questions, here we systematically study the relationship between cell fate and cell phylogeny as derived using lineage barcodes. First, we examine how phylogeny of cells can be used to understand the dynamics of the progenitor states that gave rise to them. We define quantitative fate maps that specify the fate dynamics of a progenitor field—a collection of progenitor states [20] that give rise to a set of observed cells (**Figure 1**). We then establish a generative model of cell phylogenies based on predetermined quantitative fate maps (**Figure 1A**). Using generated phylogenies, we develop a strategy, ICE-FASE, to map the order of progenitor states and quantify their commitment time, commitment bias, and population size from phylogeny, thus reconstructing the original quantitative fate map (**Figure 1B**). We find that successful quantitative fate map reconstruction requires time-scaled cell phylogeny wherein branch lengths correlate with interdivision times. It further requires adequate representation of progeny of each progenitor state among the terminal cells of the phylogenetic tree. These results demonstrate that phylogeny of a small number of cells can inform progenitor dynamics at the organism level. Second, we examine whether phylogenies inferred from lineage barcodes can reconstruct quantitative fate maps. We establish a barcoding mutagenesis model and parametrize it using experimental data from synthetic barcoding in mice. We then use the model to simulate realistic lineage barcodes based on time-scaled cell phylogenies (**Figure 1C**). To recover time-scaled phylogenies from lineage barcodes, we establish a generalizable and scalable method, Phylotime, that clusters cells based on temporal distances estimated from maximum likelihood of the mutagenesis model (**Figure 1D**). We demonstrate that Phylotime, coupled with ICE-FASE, enables quantitative fate map reconstruction from lineage barcodes (**Figure 1D,B**). Finally, we validate our methods in an experimental system with cultured stem cells grown based on preset fate parameters (**Figure 1E,D,B**). Overall, our results demonstrate how lineage barcodes from single time-point measurements can be used to decipher quantitative fate maps that capture the fate dynamics of progenitor populations long after their differentiation.

## Results

### Modeling cell phylogeny based on quantitative fate maps

To study the relationship between cell phylogeny and fate dynamics, we began by establishing the quantitative fate map, a model that specifies a developmental process wherein dividing cells assume increasingly diverse but decreasingly potent progenitor states over time before arriving at their terminal fates. Each progenitor state is defined by its potency, which is the set of terminal states it is capable of producing, and is associated with a commitment event, when its cells transition to less potent downstream states. The commitment event confers each progenitor state three additional defining parameters: i) commitment time, defined as the time when a progenitor state’ s cells commit to the downstream states, ii) population size, defined as its number of cells at commitment time, and iii) commitment bias, defined as the proportions of its population committing to each downstream state. Progenitor states ultimately give rise to terminal states, which are the states of cells at the point they are sampled or observed. In summary, a quantitative fate map defines the fate dynamics of a progenitor field—a collection of progenitor states [20] that give rise to a set of observed cells.

We next constructed a panel of 127 test quantitative fate maps covering diverse developmental scenarios (**Figure 2**). Representing increasing field sizes, the maps are in three categories of 15, 31, or 63 progenitor states, producing 16, 32 or 64 terminal cell states, respectively. We label progenitor and terminal states with “P”s and “T”s followed by numerals, respectively (**Figure 3A**). Within each category, the topologies of the maps range from perfectly balanced to highly unbalanced (**Figure 2**) as measured by the BSUM imbalance index [21]. In more unbalanced maps, progenitor states tend to split into increasingly unequal numbers of eventual terminal states (see Methods). Progenitor states in each map commit to two downstream states with prespecified ratios that were randomly drawn to cover a range of commitment biases within each map (**Figure 3A**, Methods). Commitment time for each progenitor state is also preset and randomly selected between *t* = 1. 8 to 9. 8 days subject to the topology constraints, based on the beginning of fate restrictions and the end of organogenesis in mouse development (**Figures 2 and 3A**, see Methods). Each fate map starts with a single founder cell at time *t* = 0 that divides according to the reported division rates during mouse development [22]. This cell division rate, which is the same for all progenitor states, together with their commitment biases determine the progenitor population size at different points in time (**Figure 3B, Figure S1**). Fate maps continue to *t* = 15 days when terminal cells can be sampled for observation. Supplementary Data 1 provides a complete accounting of each map. Together, these fate maps represent a broad range of complex developmental scenarios.

**Figure 2.**
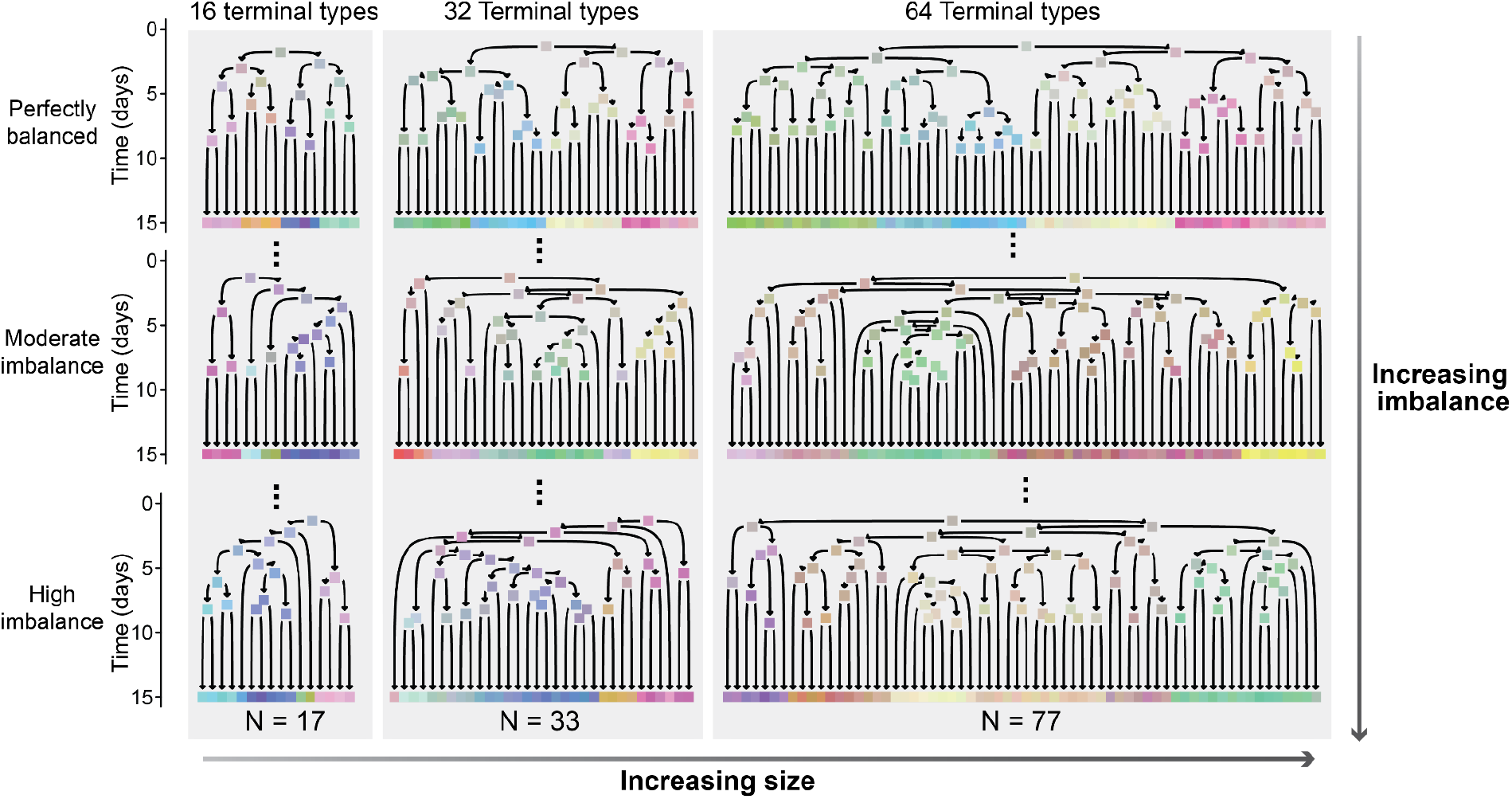
A test panel of 127 quantitative fate maps covering a broad range of developmental scenarios. The fate maps are categorized by three sizes of 16, 32, and 64 terminal cell types. Three examples from each size are shown in each column. Within each size category, topologies range from perfectly balanced (top row) to highly unbalanced (bottom row). Arrows represent cell states, colored rectangles represent their commitment events. Rectangles at day 15 represent terminal states at the time of sample collection. Each map also specifies the cell commitment bias and division rate for each progenitor state, neither of which are shown in this figure.

**Figure 3.**
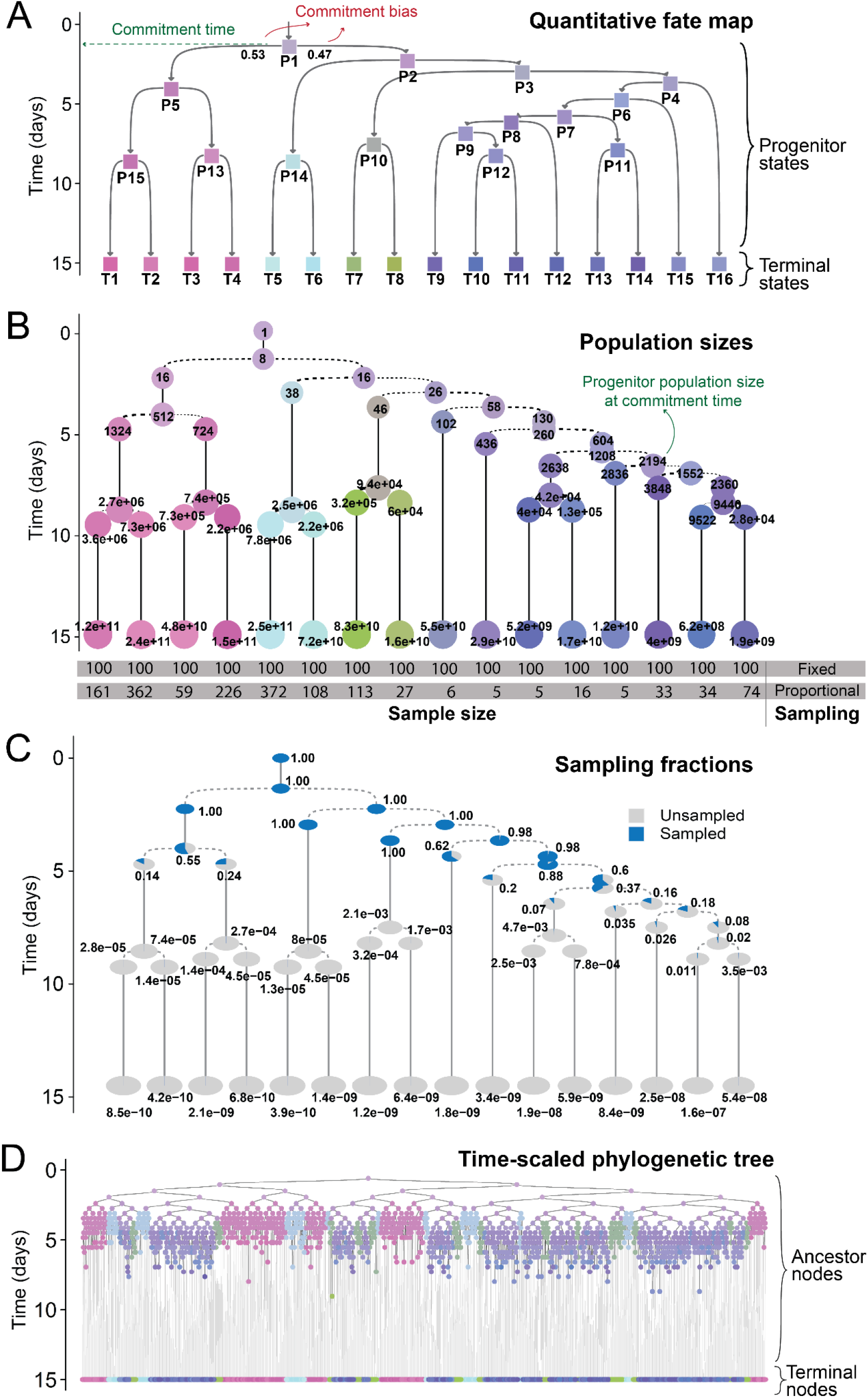
Simulating time-scaled phylogenetic trees of sampled cells based on a known quantitative fate map. (**A**) Topology of a quantitative fate map specifies commitment time and bias for each progenitor state. (**B**) Quantitative fate map is populated with cell numbers at each time point based on cell division rates; these cells are apportioned to various states based on commitment biases. A number of cells are sampled from each terminal state based on the experimental sampling strategy as shown in the gray boxes on the bottom. (**C**) Sampling of terminal states dictates the fraction of progenitor states at each time point that are ancestral to sampled cells at all points in time. (**D**) Phylogeny of sampled cells is created by merging sampled terminal cells into ancestral nodes.

We next established a generative model of cell phylogenies for terminal cells based on a quantitative fate map. Generating the entire tree of cell divisions for billions of terminal cells (**Figure 3B**) is computationally impractical. To overcome this problem, our model draws inspiration from coalescent theory in population genetics [23,24]. It starts with a subset of terminal cells that are sampled from terminal states and merges them going backward in time to create their phylogeny in three steps (see Methods). First, terminal cells are randomly sampled in each state based on either fixed or proportional sampling (**Figure 3B**). Under fixed sampling, a fixed number of cells are sampled from each terminal state, similar to experiments where target terminal cell types are first identified and then collected using sorting or other methodology. Under proportional sampling, each terminal cell state is sampled based on its share of the total population, similar to experiments where cells are sampled without prior knowledge of their state (see Methods). This step establishes the number and state of terminal nodes in the phylogenetic tree. Second, the number of progenitor cells ancestral to sampled terminal cells at all earlier times is generated. To do so, starting from the number of sampled cells of each state in the terminal point in time (*t* = 15), the number of their ancestors in the prior time point (*t* − Δ*t*) is drawn based on the total cell number at each time point. This process is repeated recursively until reaching the founder at *t* = 0 (**Figure 3C**, see Methods for details). This step establishes the number of ancestors, or internal nodes, in the phylogenetic tree at each time point. Finally, nodes at each point in time are randomly connected to their progenitors in the earlier time point to create the branches of the phylogenetic tree. At the times of commitment events, cells of less potent states are connected with the combined pool of their ancestors. This leads to terminal cells gradually merging into increasingly common ancestors according to the numbers of ancestors over time established in the previous step. This sequence of merges forms the topology of the phylogeny and the times between merges form branch lengths (**Figure 3D)**. This approach generates time-scaled phylogenies for sampled cells based on their progenitors’ fate map in a computationally efficient manner.

We applied our model to generate time-scaled phylogenies of sampled cells for each quantitative fate map in our test panel. For each map, we used both fixed and proportional sampling; for each sampling, we simulated two phylogenies. In all cases, an average of one hundred cells were sampled from each terminal state in each map. Together, these results represent 508 experiments (127 maps x 2 sampling strategies x 2 replicates) wherein phylogeny is determined for a small fraction of terminal cells derived from complex fate landscapes. With the benefit of knowing their underlying fate maps, we will use these 508 phylogenies in the ensuing sections to establish quantitative fate mapping algorithms and evaluate the reach and limitations of retrospective phylogenetic reconstruction approaches.

### Reconstructing fate map topology using time-scaled cell phylogeny

These simulated time-scaled phylogenies, in their topologies and branch lengths, embody the order and timing of cell divisions that connect sampled cells (**Figure 4A**). Going from the root of a tree towards its terminal branches, nodes become gradually restricted in their potency, as can be observed by the diversity of their terminal progenies (**Figure 4A,B**). When and which cell fate restrictions happen are clues to cell fate commitments. Therefore, to derive fate map topology from time-scaled phylogenies, we focused on putative cell fate restriction events. First, we annotated each node in the phylogenetic tree with the states of its observed terminal descendants, which we refer to as the node’ s observed fate (**Figure 4B**). Next, we compared the observed fate of each node with that of its two daughter nodes to identify nodes with both descendants having a more restricted fate (**Figure 4B**). For instance, if an internal node leads to terminal cell types T7, T8, and T9 but cells of type T7 and T8 are only seen in one of its branches and T9 only in the other, this node implies divergence of the terminal cell types T7 and T8 from T9. We refer to these nodes as observed FAte SEparation (FASE) between respective terminal fates. The prior example constitutes a T7–T9 and a T8–T9 FASE. The temporal distribution of FASEs that connect a pair of terminal cell types provides a measure of their developmental distance: cell types whose lineages separated early in development would be connected by FASEs that are closer to the root in the phylogenetic tree whereas those whose lineages separated later in development would be connected by FASEs that are closer to terminal branches (**Figure 4C**). Therefore, we defined the distance between terminal cell fates T1 and T2 as the mean temporal distance of T1–T2 FASEs to their terminal cells and compiled a pairwise distance matrix between all terminal cell types (**Figure 4D**, see Methods). Applying a distance-based clustering method (UPGMA) to this matrix, we obtained the topology of the fate map (**Figure 4E**). This fate map topology connects observed terminal cell states through a hierarchy of inferred progenitor states, each with the potency to give rise to two or more terminal cell types. We label these inferred progenitor states by “*iP”* followed by a numeral. To summarize, this strategy, which we call the FASE algorithm, produces a hierarchy of “inferred” progenitor states based on specific patterns of potency that result in the observed terminal states.

**Figure 4.**
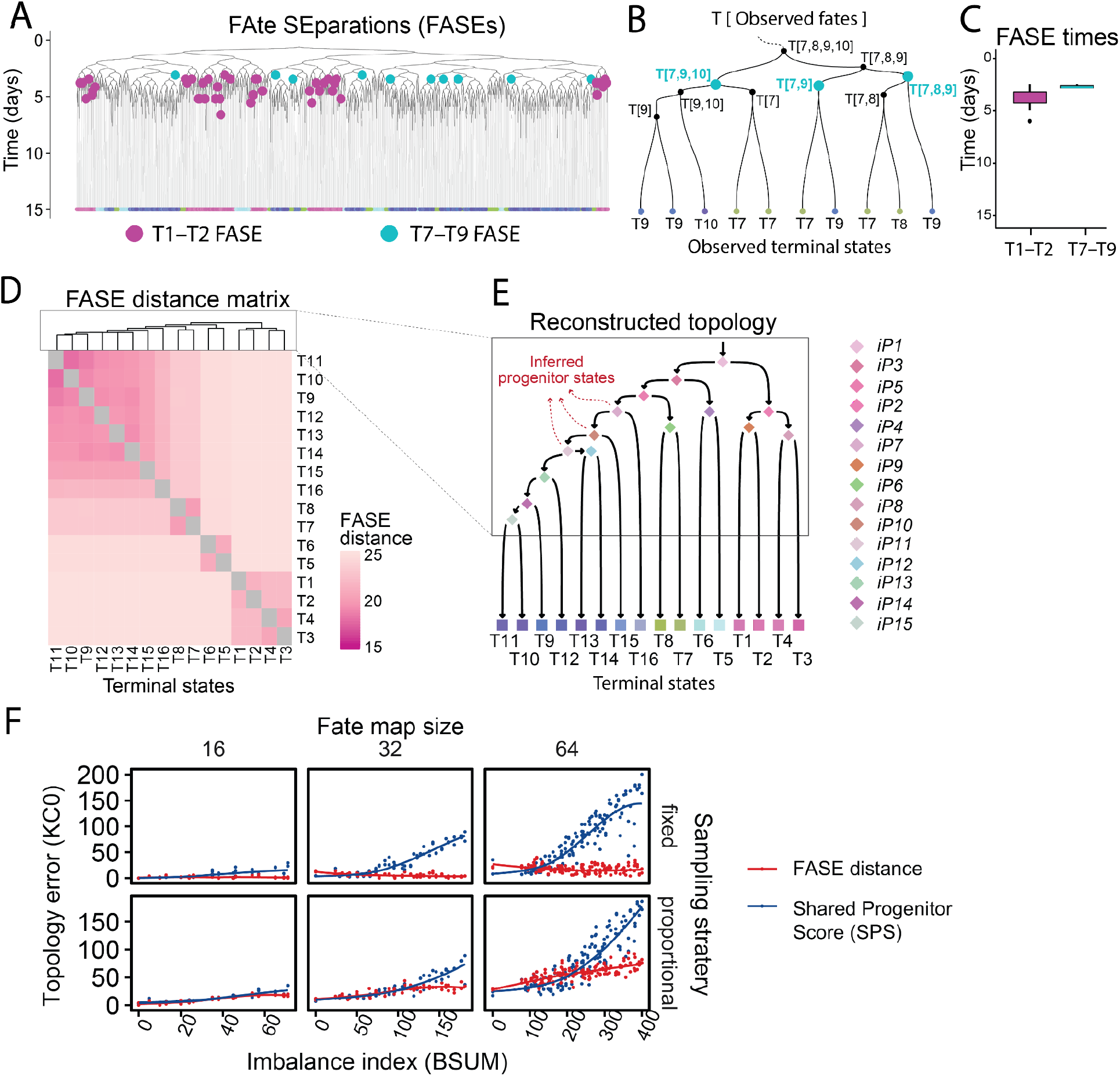
Reconstructing fate map topology from time-scaled phylogeny of sampled terminal cells. (**A**) A time-scaled phylogenetic tree of sampled terminal cells. Terminal cells of different states have been labeled by different colors. FAte SEparation (FASE) events between terminal states T1 and T2 (purple) as well as those between T7 and T9 (teal) have been labeled as examples. (**B**) A zoomed-in section of the tree in panel A where each internal node is labeled by its observed fate and T7–T9 FASEs have been labeled by teal circles. (**C**) Boxplot showing the distribution of T1–T2 and T7–T9 FASE times in the tree in A, from which the distance between terminal states was derived. (**D**) Heatmap showing the FASE distance matrix for all pairs of terminal states in the phylogenetic tree in panel A. Dendrogram on top shows the hierarchical clustering results of the heatmap. (**E**) The fate map topology reconstructed by hierarchical clustering of the FASE distance matrix. Squares denote observed terminal states, diamonds denote inferred progenitor states. (**F**) Scatter plots showing the distance between fate map topologies inferred from the time-scaled phylogenetic tree and the true fate map topology in all 508 simulated phylogenies. The algorithm using FASE times for topology inference is compared to one using the Shared Progenitor Score (SPS) as a measure of similarity. Results are broken down in different plots by the total number of terminal states, the experimental sampling strategy, and by fate map imbalance on the x-axes. Solid lines show trend lines obtained through locally weighted smoothing (LOESS).

We applied the FASE algorithm to reconstruct fate map topology for each simulated phylogeny in our panel. For comparison, we also used the shared progenitor score (SPS) [10]. In each case, we compared the reconstructed topology to its corresponding true fate map using the Kendall-Colijin (KC) metric with its tuning parameter (λ) set to zero (KC0) [25]. A KC0 distance of zero indicates identical topologies; KC0 distances larger than zero indicate increasing differences between topologies. The results show that our FASE strategy faithfully reconstructs fate map topology in almost all tested phylogenies, outperforming SPS across the board (**Figure 4F**). Under fixed sampling, the FASE algorithm predicts perfectly accurate topologies in 16 and 32-terminal cell state fate maps irrespective of imbalance. It only shows modest decrease in accuracy with high degrees of imbalance under proportional sampling (**Figure 4F**). Moreover, fixed sampling outperforms proportional sampling, likely because it ensures better representation of small terminal populations. This result suggests that a fixed sampling strategy is more robust in lineage topology reconstruction using barcoding approaches than a proportional sampling strategy. Taken together, these results establish a robust and scalable method to reconstruct fate map topology from the phylogeny of sampled cells that scales to complex fields.

### Quantitative characterization of progenitor states using cell phylogeny

The fate map topology derived from the lineage of sampled cells effectively identifies a series of inferred progenitor states (*iPs*) each with a distinct potency and fate restriction pattern (**Figure 4E**). As the internal nodes of time-scaled phylogenies correspond to cells in these progenitor states, we sought to use them to further characterize the dynamics of these progenitor states at the cell population level. We therefore assigned each internal node in the phylogenetic tree to an inferred progenitor state or a terminal state based on fate map topology: given the observed fate of the internal node, we assigned it to the least potent progenitor state that contains the node’ s observed fate (**Figure 5A**). For example, a node with an observed fate of [T11, T13, T14] can be assigned a more potent inferred state (*iP11*) capable of [T9, T10, T11, T12, T13, T14] if the now-reconstructed topology of the fate map indicates that *iP11* differentiates into fates [T13, 14] and [T9, T10, T11, T12] (**Figure 5B**). To assess the fidelity of these assignments, we compared the inferred states of internal nodes in all phylogenies to their true states which are known, as these phylogenies were simulated. Where derived fate map topology differed from the truth, only correctly resolved progenitor states were considered. A progenitor state is correctly resolved if there exists a corresponding state in the true fate map with the same potency and commitment pattern (**Figure 5C**). The only error was assigning an internal node to a progenitor state less potent than its true state, which occurred, on average, for 21% of all nodes in each time-scaled phylogeny among the 508 experiments (**Figure 5C,D**). These errors are caused by failures to sample all the different terminal states a progenitor cell has led to and emerges when the number of sampled cells is small relative to the terminal population (**Figure 5E**). Since the sampling fraction of each progenitor state is known in simulated experiments (**Figure 3C**), we further examined its effect on resolving progenitor states. We found that a great majority of progenitor states in our panel were correctly resolved if more than 25% of their population at the time of commitment were represented among sampled terminal cells. Conversely, only a minority of progenitor states with less than a 25% sampling fraction were correctly resolved. Going forward, we will distinguish progenitor states with more than 25% sampling as adequately-sampled and the rest as under-sampled.

**Figure 5.**
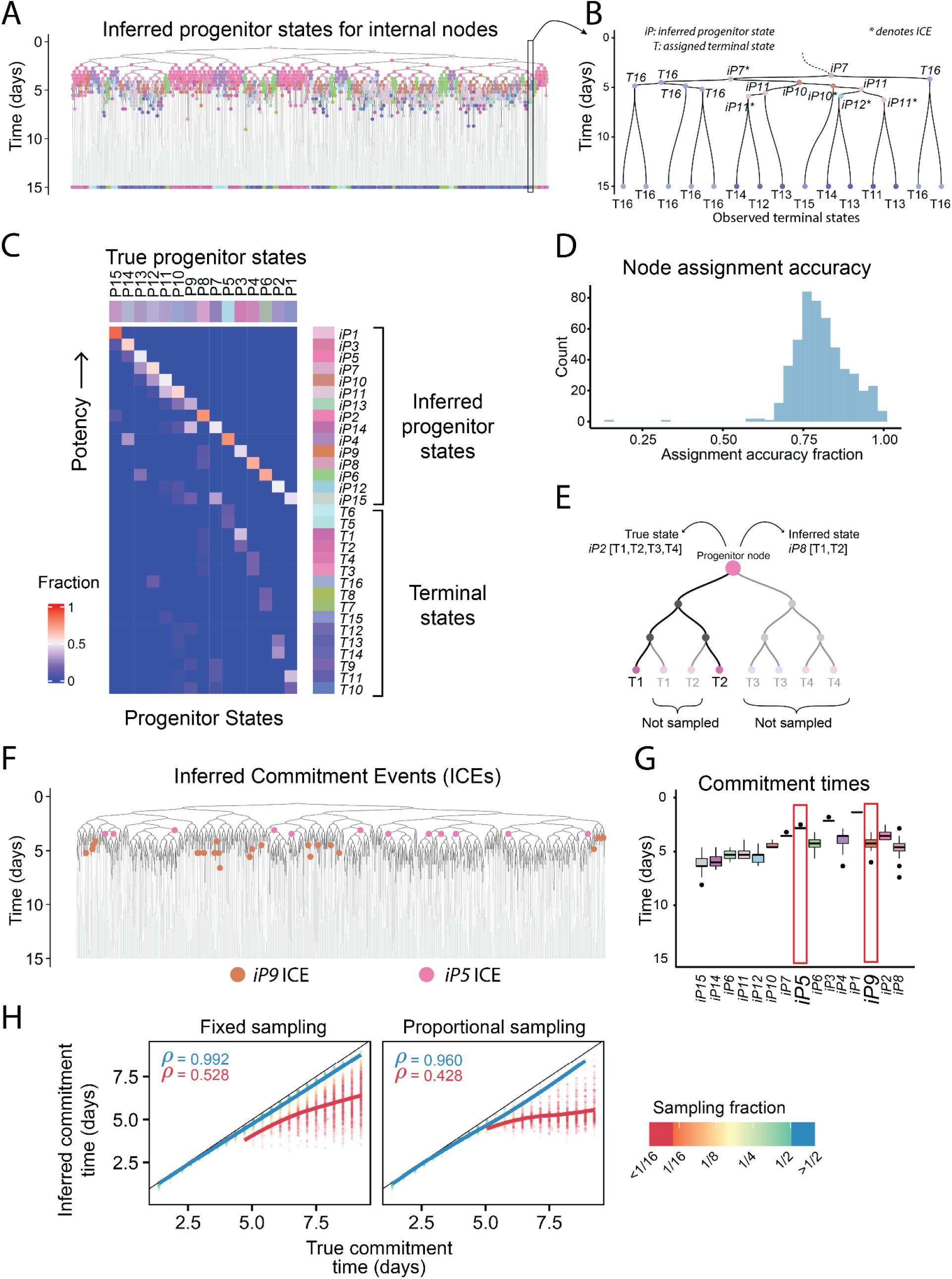
Obtaining progenitor state commitment times from phylogenies of sampled cells. (**A**) The time-scaled phylogenetic tree in Figure 4A where internal nodes are colored according to their inferred progenitor state. (**B**) A zoomed-in section of the tree in panel A where each internal node is labeled by its inferred progenitor state or terminal state. Asterisks signify ICE nodes. (**C**) Heatmap showing the agreement between inferred states and true states of all internal nodes in the tree shown in panel A. (**D**) Histogram showing the average accuracy of progenitor state assignment to internal nodes of the phylogenetic tree across the panel of 518 phylogenetic trees. On average, 79% of nodes were correctly assigned to a progenitor state, the rest were assigned to states with less potency. (**E**) Schematic showing how undersampling can lead to a node being assigned a progenitor state with less potency than its true state. (**F**) The tree in panel A where nodes corresponding to Inferred Commitment Events (ICEs) for inferred progenitor states iP9 and iP5 are shown in brown and pink, respectively. (**G**) Distribution of ICE times for all inferred progenitor states in the tree from panel A, representing each progenitor state’ s commitment time. The ICE times for *iP5* and *iP9* are boxed in red. (**H**) Scatterplots showing the correlation between actual commitment time of each progenitor state to the value inferred from the phylogenetic tree across all 508 simulated phylogenies broken down by experimental sampling strategy. Dots are colored based on progenitor sampling fraction and according to the key on the right. Trendlines (LOESS) for adequately-sampled and undersampled progenitor states are shown in blue and red, respectively. *ρ* indicates Spearman’ s correlation coefficient. The blue value corresponds to progenitors with sampling fraction better than 25%; the red value to those with sampling fraction equal to or less than 25%.

To derive the commitment time of each progenitor state, we focused on when its inferred nodes transition to less potent states by defining a set of Inferred Commitment Events (ICEs). An ICE is a node whose inferred state is more potent than both of its descendants (**Figure 5B,F**). For example, in Figure 5B, when an internal node is assigned to *iP7* (capable of T9 through T16) and splits into nodes with assigned states of *T16* and *iP10*, we count this node as an ICE for *iP7*. The temporal distribution of each progenitor state’ s ICEs indicate when the progenitor commits to its downstream fates (**Figure 5G**). Unlike FASEs, which are defined for each pair of terminal fates, ICEs are defined with respect to inferred progenitor states. If a node is an ICE, it is also a FASE for at least some pair of terminal states, but the opposite is not necessarily true. ICE uses information from the fate map topology to identify a more confident set of nodes that represent state transitions. We thus defined the commitment time for a progenitor state as the mean of its ICE times (**Figure 5G**). Across all progenitor states, these ICE times captured the relative timing of commitment events as indicated by a high rank correlation (Spearman’ s ρ=0.849 for fixed sampling and ρ=0.901 for proportional sampling for progenitor states that are correctly resolved in fate map topology) (**Figure 5H**). However, across progenitor states that were adequately sampled, the ICE times captured the exact timing of commitments as indicated by a low root mean square error (RMSE=0.298 days for fixed sampling and RMSE=0.282 days for proportional sampling). Fixed and proportional sampling performed comparably, though fixed sampling performs better for under-sampled progenitors. These results establish ICE times as estimates for the commitment times of the progenitor states from time-scaled phylogenies of sampled cells.

We next assessed whether population size and commitment bias of a progenitor state can be obtained from time-scaled phylogenies. We define progenitor population size as the number of cells in a progenitor population immediately before they commit to their downstream fates and commitment bias as the ratio of these cells that commit to each downstream fate. To estimate population size for a progenitor state, we identified the subset of all branches in the phylogenetic tree that, first, are present at the progenitor state’ s inferred commitment time and, second, connect nodes assigned as either the progenitor state itself or any of its upstream or downstream states (**Figure 6A,B**, see Methods). These branches represent cells of the progenitor state that are present at its time of commitment. We then counted the number of incoming nodes to these branches as the population size (**Figure 6C**). For commitment bias, we calculated the ratio of branches that end in each of the downstream fates irrespective of their parental state (**Figure 6D**, see Methods). Applying this algorithm to our panel of time-scaled phylogenies, we found that the ability to estimate population size and commitment bias for a progenitor state depend heavily on its sampling fraction as well as the sampling method.

**Figure 6.**
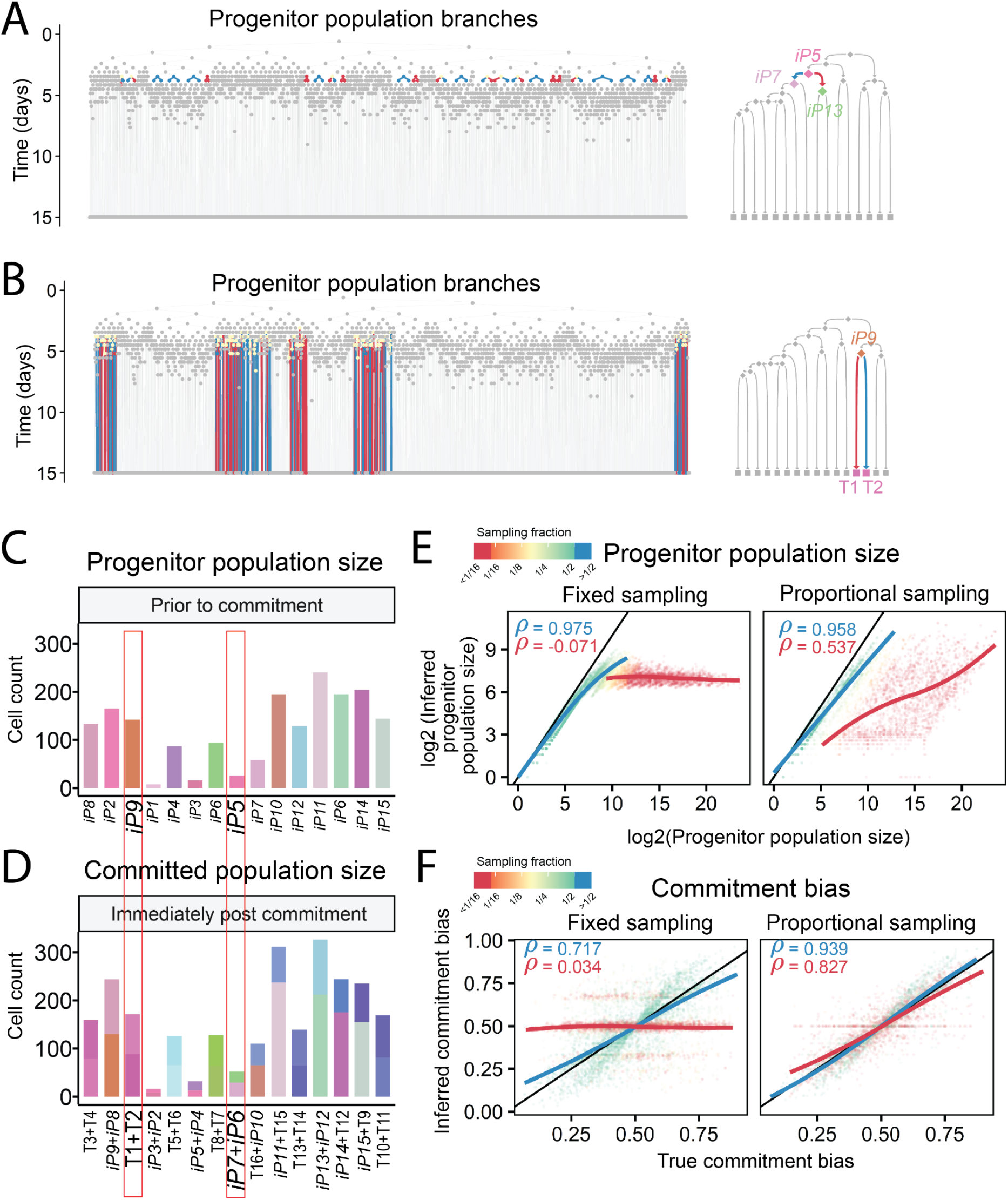
Obtaining progenitor population size and commitment bias from phylogeny of sampled cells. (**A**) The time-scaled phylogenetic tree from Figure 4A is shown and branches corresponding to inferred progenitor state *iP5* that are present at its commitment time are colored based on the state of the node they end in according to the key shown on the fate map to the right. (**B**) Same as A, but for *iP9*. (**C**) Barplots showing the calculated population size of each progenitor state in A based on the nodes before its branches at its commitment time. (**D**) Barplots showing the calculated post-commitment population size of each progenitor state in A immediately after its commitment time based on the nodes after its branches at its commitment time and stacked according to the downstream state they lead to. (**E**) Scatterplots showing the correlation between true population size of each progenitor state to the value inferred from the phylogenetic tree across all 538 simulated phylogenies broken down by experimental sampling strategy. Dots are colored based on progenitor sampling fraction and according to the key on the right. Trendlines (LOESS) for adequately-sampled and undersampled progenitor states are shown in blue and red, respectively. *ρ* indicates Spearman’ s correlation coefficient. (**F**) Scatterplots showing the correlation between actual commitment bias of each progenitor state to the value inferred from the phylogenetic tree across all 538 simulated phylogenies broken down by experimental sampling strategy. Dots are colored based on progenitor sampling fraction and according to the key on the right. Trendlines (LOESS) for adequately-sampled and undersampled progenitor states are shown in blue and red, respectively. *ρ* indicates Spearman’ s correlation coefficient.

Population size estimates of adequately-sampled progenitor states agree well with their actual size (Spearman’ s ρ=0.975 for fixed and ρ=0.958 for proportional sampling) (**Figure 6E**). For undersampled progenitor states, on the other hand, population size estimates are capped at the number of their sampled terminal progeny, thus performing poorly for the proportional sampling scheme (Spearman’ s ρ=0.537) and being completely non-informative for the fixed sampling scheme (Spearman’ s ρ=-0.07) (**Figure 6E**). Commitment bias estimates showed a different behavior: proportional sampling allowed for estimation with a minor effect from progenitor’ s sampling fraction (Spearman’ s ρ=0.939 and ρ=0.827 for adequately and undersampled progenitor states, respectively) (**Figure 6F**). Fixed sampling strategy, on the other hand, produced reasonable estimates only for adequately-sampled progenitor states’ commitment biases (Spearman’ s ρ=0.717) and was uninformative for undersampled progenitor states (Spearman’ s ρ=0.034) (**Figure 6F**). These observations demonstrate the importance of sampling fraction for accurate characterization of progenitor states. They further indicate more effective estimation of population size and commitment bias with proportional sampling schemes, ostensibly due to the correlation between the size of terminal populations and that of their progenitor. Together, these results establish a strategy for estimating progenitor population size and commitment bias from time-scaled phylogenetic trees and define critical parameters for robust estimation.

### Robustness of phylogeny-based quantitative fate map estimates

So far, we have established the central role of progenitor state sampling fraction as an indicator of the ability to infer quantitative fate map parameters. However, this parameter cannot be directly observed from cell phylogeny or other data collected from sampled terminal cells (**Figure 3C**). Hence, it is important to find a proxy for the true sampling fraction of each progenitor state that can be derived from sampled cells alone. Here, we introduce one such proxy: estimated progenitor state coverage (PScov), which is defined as the terminal sample size of a progenitor state divided by its estimated progenitor population size. By terminal sample size, we refer to the sum of the number of sampled cells of all the terminal states that the progenitor state is capable of. Intuitively, this statistic indicates how many terminal descendents are being sampled per each progenitor cell. We found that a high PScov is indeed predictive of high sampling fraction, auROC = 0.969 (CI: 0.967 - 0.971) (**Figure S2A**). For example, the majority of states that are sampled more than 25% also have a PScov larger than 2.5 (**Figure S2B**). Therefore, this estimated progenitor state coverage makes it possible to assess the robustness of quantitative fate map parameters for each progenitor state based solely on the phylogeny of terminal cells.

### Modeling and simulating lineage barcoding in development

The results above indicate that time-scaled phylogenies of sampled cells can identify progenitor states of common fate and their dynamics. The time-scaled phylogenies that were used thus far were known—representing the exact sequence and timing of events as simulated. In actual experiments, phylogeny must be inferred from lineage barcodes. However, such inferred phylogenetic trees are inherently error-prone for two reasons. First, exhaustive phylogenetic tree search to guarantee optimality is not computationally practical for hundreds of terminal cells, and practical heuristic algorithms do not guarantee optimality [26] despite the recent advances in distributed computing [27]. Second, barcoding strategies employ a limited number of mutation sites which encode a limited amount of information; how close any inferred tree—including the optimal tree—is to its true phylogeny remains uncertain. We therefore asked: can phylogenies inferred from lineage barcodes be accurate enough to recreate quantitative fate maps despite their errors? To address this question, in this and the ensuing two sections, we will: i) describe a model to generate realistic barcoding outcomes in cells, ii) establish a scalable strategy for inferring time-scaled phylogenies from barcodes, and iii) evaluate the feasibility of quantitative fate mapping using barcode-inferred phylogenies.

First, we established a general mutagenesis model. The model comprises barcoding sites that are present in each cell and inherited to its daughters (**Figure 7A**). Barcoding sites are unmutated in the founder cell at the beginning (i.e., *t* = 0) and once activated, start accumulating heritable mutations over time (**Figure 7A**,**B**). Each site mutates independently from other sites in the system according to a Poisson point process with a constant rate after activation. Each mutation event converts an unmutated active copy of the site into one of many possible mutated inactive alleles, each with a distinct emergence probability and unable to mutate further (**Figure 7C,D** and **Methods**). Therefore, the parameters of the barcoding model are the number of sites, their mutation rates, and each site’ s mutant allele emergence probabilities.

**Figure 7.**
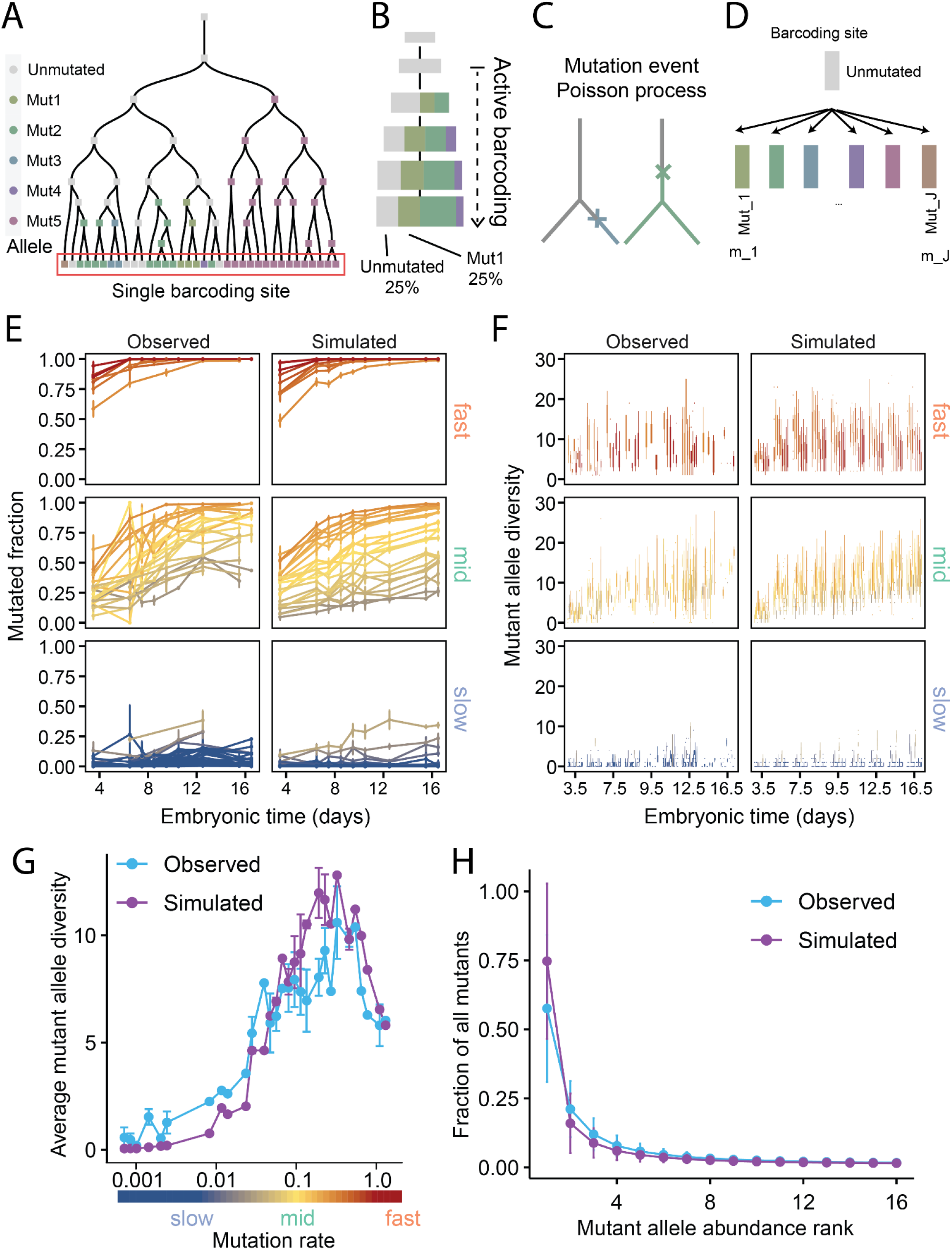
Lineage barcoding model and agreement of simulated barcoding outcomes to those observed in MARC1 mice. (**A**) A phylogenetic tree with a single barcoding element shown as a box and its allelic state denoted with color according to the key on the right. The barcoding element is inherited from each cell by its daughters. Only the unmutated allele (gray) can change, mutated alleles (colored), once formed, are inherited from a cell by all its daughters. (**B**) Accumulation of mutations over time upon barcoding activation across a population of cells. (**C**) Mutation events (crosses) take place according to a Poisson point process. (**D**) Mutation events convert the unmutated allele to one of many possible mutated alleles (*Mut*_1_,…, *Mut*_*J*_) (Mut_1_ to Mut_j_), each with an emergence probability (m_1_ to m_j_). (**E**) Line-plots showing the mutated fraction of MARC1 barcoding elements over time in observed (left) and simulated (right) embryos for fast, mid, and slow categories of mutation rates. Means ± SEM are shown. Simulations and experimental observations for the same element are shown in the same color. (**F**) Boxplots showing the number of mutant alleles for each MARC1 barcoding element over time in observed (left) and simulated (right) embryos for fast, mid, and slow categories of mutation rates. Simulations and experimental observations for the same element are shown in the same color. (**G**) Line-plots showing the average mutant allele diversity as a function of barcoding element mutation rate for observed (blue) and simulated (purple) embryos. Means ± SEM are shown. (**H**) Line-plots showing the prevalence of a mutant allele among all mutant alleles as a function of its rank in observed (blue) and simulated (purple) embryos. For example, a rank of one denotes the most abundant allele. Means ± SE are shown.

This model is broadly applicable to synthetic and natural barcoding systems; here, we parameterize it based on the MARC1 (Mouse for Actively Recording Cells!) system [9] wherein extensive embryonic barcoding data are available [28]. In MARC1 mice, somatic mutations are induced in tens of independent homing guide RNA loci (hgRNAs) [29] starting at the 2-cell stage. We estimated the mutation rates of MARC1 hgRNAs (i.e., λ, rate of the Poisson process) using embryonic time course data [28] (**Figure S3A**,**B**, see Methods). We estimated emergence probabilities of mutant alleles for each hgRNA by adapting the inDelphi algorithm that predicts CRISPR-Cas9 mutations [30]. We compared and verified inDelphi’ s predictions against published MARC1 data [28] (**Figure S3B**, Supplementary Data 2, see Methods). To test this MARC1 barcoding model, we simulated barcoding in whole-mouse embryos for E3.5 to E16.5 in samples of 2,000 cells (or fewer when there were fewer than 2,000 cells in the organism) and compared the results to that of experiments (see **Methods**). Overall, we observed broad agreement between experimental and simulated barcoding results (**Figure 7E–H**). First, the distribution of mutated fractions over the course of embryogenesis agrees between simulated and experimental results for hgRNAs with a range of mutation rates (**Figure 7E**). Second, the total number of distinct mutant alleles (i.e. the mutant allele diversity) during embryogenesis were consistent between experiments and simulations (**Figure 7F**). Third, in both systems, as the hgRNA mutation rates increase, the diversity of mutated hgRNA alleles increase because there are more mutagenesis events; however, at the fastest mutation rates, barcoding sites reach 100% mutated when there are fewer total cells, and the total diversities drop as a result (**Figure 7G**). Fourth, the composition of mutant alleles within embryos agree: for both simulated and observed embryos, after ranking all mutant alleles based on their frequencies, alleles of similar rank account for similar percentages of the mutated cells (**Figure 7H**). Taken together, these results suggest that our simulation is comparable to actual lineage barcoding experiments and produces realistic barcoding results.

Finally, based on the above stochastic barcoding model and the MARC1 system’ s parameters, we simulated mutagenesis in our panel of 508 phylogenies assuming 100 hgRNA per cell (see Methods). The results are 508 simulated barcoding experiments where, similar to real-life experiments, the barcode and terminal state information is known for the sampled panel of single cells (**Figure 8A**; **Supplementary Data 4**).

**Figure 8.**
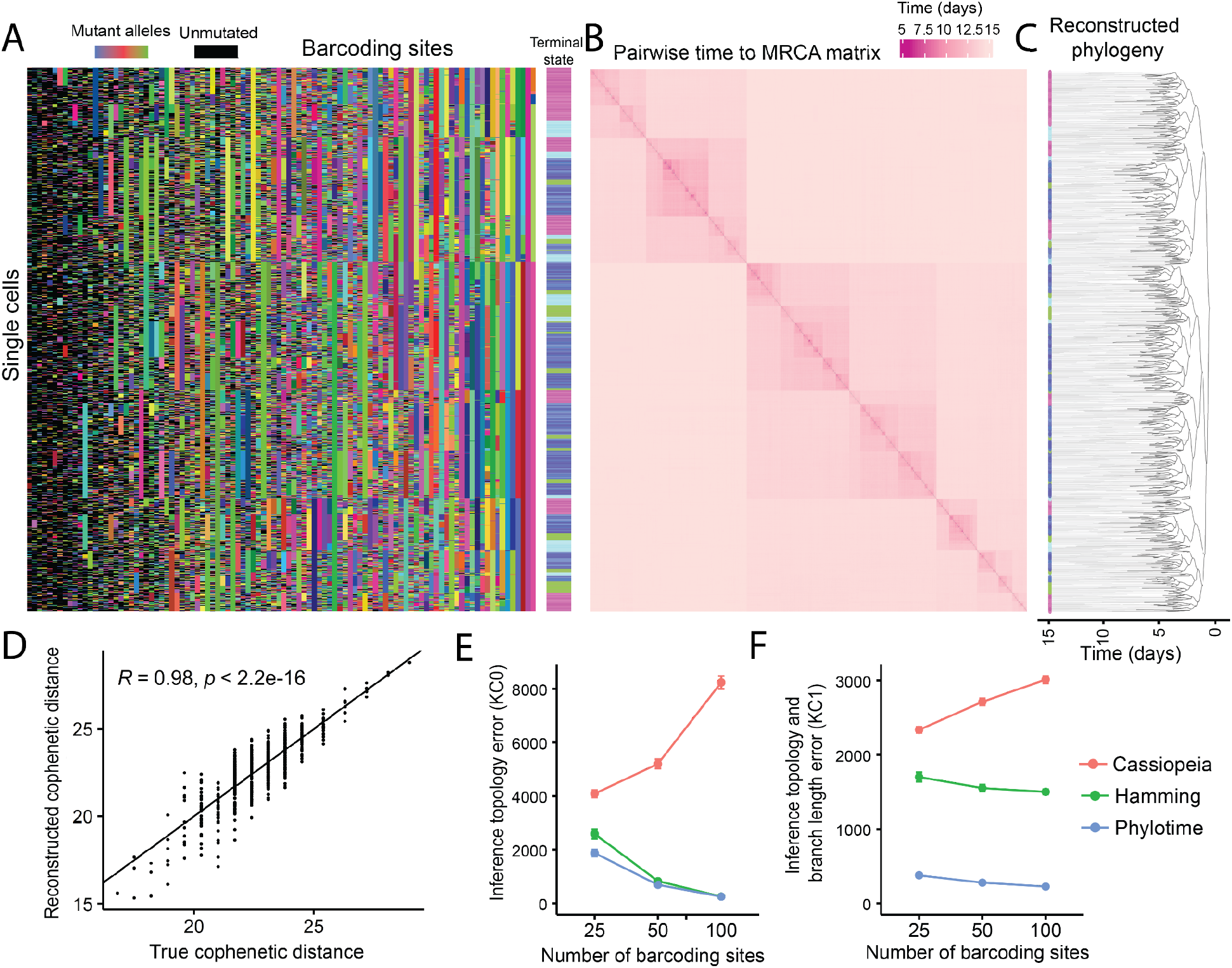
Accurate reconstruction of time-scaled phylogenetic trees using Phylotime. (**A**) The output of a simulated barcoding experiment shown as a barcode matrix with each hgRNA barcoding element as a column and each cell as a row. Colors correspond to mutant alleles; black is an unmutated allele. The color bar on the right shows the state of each terminal cell. (**B**) Heatmap showing the pairwise time to most recent common ancestor (MRCA) for all cells in A. (**C**) Time-scaled phylogenetic tree reconstructed by applying a clustering algorithm to the matrix in B. Colors on the terminal branches signify the observed state of the cell. (**D**) Scatter plot showing the correlation between the Phylotime-inferred and true cophenetic distances between all pairs of cells in A. Trendline is shown. (**E**) Error of phylogenetic reconstruction using Phylotime, Hamming distance with UPGMA, and Cassiopeia, with 25, 50, or 100 barcoding elements when considering only tree topology (KC0 distance) across the panel of 518 simulated barcoding experiments. Means ± SEM are shown (N=518). (**F**) Error of phylogenetic reconstruction using Phylotime, Hamming distance with UPGMA, and Cassiopeia, using 25, 50, or 100 barcoding elements when considering both tree topology and branch length. Cassiopeia phylogenies were scaled to the same total time as the reference trees. Means ± SEM are shown (N=518).

### Reconstructing time-scaled phylogenies from single-cell lineage barcodes

To infer quantitative fate maps in each simulated barcoding experiment, single-cell barcodes must be converted to a time-scaled phylogenetic tree. However, many current methods for phylogenetic reconstruction based on lineage barcodes lack a mutagenesis model specific to lineage barcoding. As a result, their inferred phylogram branch lengths are often in arbitrary units. Those strategies that do involve a barcoding mutagenesis model [31] depend on optimization techniques that are not scalable to thousands of sampled cells as we have simulated in each experiment. To address this gap, we developed a method to infer phylogenies with branch lengths measured in actual time units that readily scales to thousands of cells. We first compute a maximum likelihood estimate of the time that separates a pair of cells from their most recent common ancestor (time to MRCA) for all pairs of terminal cells (**Figure 8B**, see Methods). We apply UPGMA hierarchical clustering [32] to the pairwise temporal distance matrix to obtain a time phylogenetic tree (**Figure 8C**). We call this approach, which scales in polynomial time, PHYlogeny reconstruction using Likelihood Of TIME (Phylotime).

To evaluate Phylotime’ s performance in reconstructing time-scaled phylogenies, we first compared MRCA times estimated in our simulated barcoding experiments to that derived from the corresponding true trees (**Figure 8D**) and found that the two are highly correlated (Pearson’ s R = 0.98). We then applied Phylotime to all 508 simulated barcoding experiments to obtain inferred-phylogenetic trees (**Figure 8C**; **Supplementary Data 4** for all trees) and compared the topology and branch lengths of the Phylotime-reconstructed trees to the true trees, using KC0 distance for topology and KC1 distance for combined topology and branch length (**Figure 8E,F**). KC1 is the Kendall-Colijin metric weighted by branch length by setting its parameter to one [25]. A KC1 distance of zero between two trees indicates identical topology and branch lengths and distances larger than zero indicate increasing differences in branch length and topology. The results show that Phylotime’ s solutions converge to the true phylogeny with an increasing number of barcoding sites(**Figure 8E,F**) and produce time-scaled phylogenies that are by far the most accurate according to KC1 error (**Figure 8F**). Only a Hamming distance based method and Cassiopeia [33], a heuristic approach based on maximum parsimony, were compared because other common methods do not scale to this number of terminal cells and barcoding sites. Expectedly, given their scale, none of the trees were perfectly reconstructed compared to the truth, recapitulating errors that are inherent to tree inference. Despite this, Phylotime’ s results provide a test panel of inferred time-scaled phylogenetic trees, with generally accurate topology and branch length for fate map reconstruction.

### Quantitative fate map inference based on lineage barcodes

We next assessed if time-scaled phylogenetic trees inferred from lineage barcodes can faithfully reproduce quantitative fate maps despite their inherent uncertainties. We applied the ICE-FASE algorithm to all 508 time-scaled phylogenetic trees that were inferred using Phylotime from simulated barcoding experiments to derive quantitative fate maps. We then compared various parameters of these fate maps against those obtained by applying ICE-FASE to the true phylogeny. We found that inferred phylogenies perform only marginally worse than true phylogenies at estimating fate map topology, regardless of map complexity and imbalance, or the sampling strategy (**Figure 9A**). Similarly, for commitment times, population sizes, and commitment biases of progenitor states that were properly identified in the reconstructed topologies, we found that Phylotime-inferred phylogenies performed slightly worse than true phylogenies (**Figure 9B–D**). Taken together, these results show that the ICE-FASE algorithm can faithfully reconstruct quantitative fate maps and infer parameters of the progenitor states using barcode-inferred phylogenies. This finding indicates that quantitative fate mapping is feasible despite errors inherent to phylogenetic reconstruction. More broadly, these results show that quantitative fate mapping may be accomplished with current lineage barcoding technologies.

**Figure 9.**
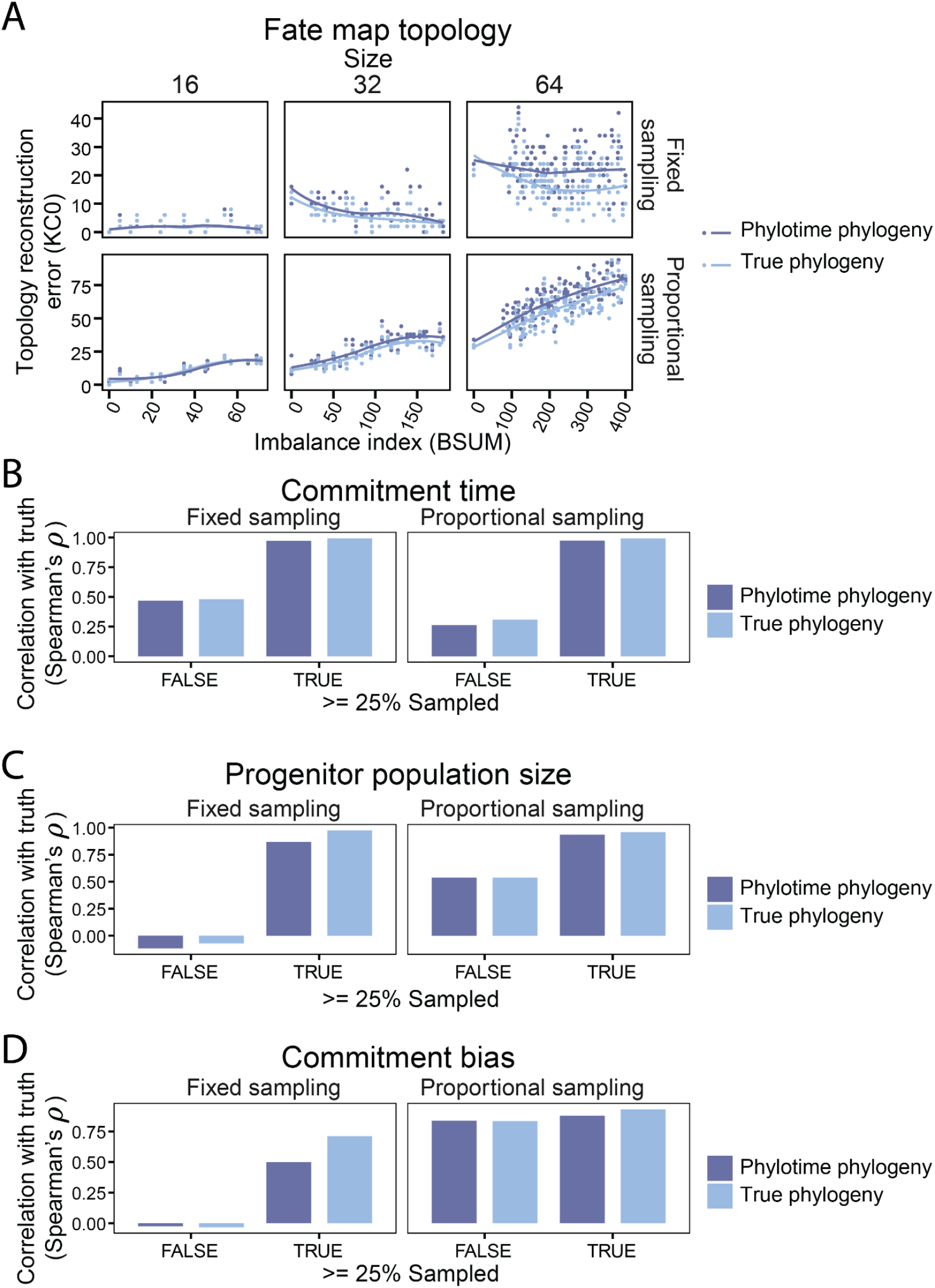
Successful quantitative fate mapping using barcode-reconstructed time-scaled phylogenetic trees. (**A**) Scatter plots showing fate map topology reconstruction error as a function of its imbalance in all the fate maps described in Figure 1 broken down by fate map size and experimental sampling strategy. Trendlines (LOESS) are shown. (**B**) Barplots showing Spearman’ s correlation between true commitment time and inferred commitment time of all progenitor states in 508 simulated experiments for progenitor states that were adequately sampled (TRUE) and undersampled (FALSE), broken down by fate map size and experimental sampling strategy. Results using true simulated phylogenies (light blue) or Phylotime-inferred phylogenies (dark blue) are shown. (**C**) Barplots showing Spearman’ s correlation between true population size and inferred population size of all progenitor states in 508 simulated experiments for progenitor states that were adequately sampled (TRUE) and undersampled (FALSE), broken down by fate map size and experimental sampling strategy. Results using true simulated phylogenies (light blue) or Phylotime-inferred phylogenies (dark blue) are shown. (**D**) Barplots showing Spearman’ s the correlation between true commitment bias and inferred commitment bias of all progenitor states in 508 simulated experiments for progenitor states that were adequately sampled (TRUE) and undersampled (FALSE), broken down by fate map size and experimental sampling strategy. Results using true simulated phylogenies (light blue) or Phylotime-inferred phylogenies (dark blue) are shown.

### In vitro validation of quantitative fate map inference using lineage barcodes

The above results prove in principle that lineage barcodes from a subset of terminal single cells can be used to retrospectively reconstruct quantitative fate maps of their progenitors. These quantitative fate maps not only define progenitor states based on their fate but also provide information about the times they were present, their numbers, and their commitment biases. This quantitative fate map reconstruction strategy constitutes a new method in the developmental biology toolbox to assess developmental systems, such as mammals, wherein multipotent progenitor populations dynamically commit to downstream fates. However, in silico models inevitably make simplifying assumptions. Our models have assumed that cells proliferate at fixed intervals with constant rate and without death, or that all the lineage barcodes can be measured without error. Therefore, we sought to validate our quantitative fate mapping strategy on an in vitro experimental model. However, there is no known biological system where these quantitative parameters for a set of progenitor states are established, making it difficult to experimentally test this novel approach. To address this gap, we created an experimental system in which quantitative fate map parameters can be controlled and then interrogated using lineage barcodes. We established a clonal human induced pluripotent stem cell (iPSC) line with 32 hgRNA barcoding sites distributed in its genome as a non-tandem array (**Table S1**). The line also includes doxycycline inducible Cas9 expression for barcoding activation (see Methods). 24 of the 32 hgRNAs were active and accumulated random mutations upon doxycycline induction (**Figure S4**).

Using this barcoding cell line, we designed a growing and splitting scheme in culture that mimics the cell fate commitment hierarchies that we have so far only simulated in silico (**Figure 10A**). In two experiments, starting from single cells, we initiated barcoding and passaged the growing cell population into an increasing number of branches at known times, numbers, and split ratios (**Figure 10B, Figures S5**,**S6**, see **Methods**). The experiments were similar except that in one experiment (E1), progenitor state #3 (P3) was split two days before progenitor state 4 (P4), whereas in the other (E2), P4 was split two days before P3. In effect, these experiments represent quantitative fate maps with the last set of cultured populations as the terminal cells and their prior ancestors as the progenitor populations. Finally, we sequenced barcodes from 192 single cells in each terminal population (see **Methods**). After data processing and filtering out low quality cells, we obtained on average 158 cells per terminal population (**Figure S7A**,**B**). E1 and E2 had medians of 27 and 26 hgRNAs detected per cell, respectively (**Figure 10C**). We imputed the alleles of undetected hgRNAs using the machine learning software “xgboost” [34] (see **Methods**) As a reference, we conducted parallel simulations on the two fate maps that represent the ground truths, with cell division rates derived from the progenitor population sizes at each split. The lineage barcodes are simulated with hgRNA mutation rates and allele emergence probabilities obtained from time-course measurements and inDelphi predictions, respectively. We then applied Phylotime and ICE-FASE to both simulated and experimental data. The experimental data reconstructed the topology correctly in both E1 and E2 (**Figure 10D**), and so we refer to the inferred states by their true state names hereafter. The PScov ranged from 1.68 to 2.36 in all non-founder (P5) progenitor states, indicating that the progenitor states were generally undersampled in these experiments. Nevertheless, in addition to the correct topology, the inferred fate maps recovered the correct orders of fate commitment in both experiments (**Figure 10D**). Moreover, the inferred maps successfully recovered the difference between E1 and E2, which was the relative order of commitment for P3 and P4 progenitor states (**Figure 10D**). This result suggests that our strategy can identify specific quantitative fate map differences in different systems. However, due to undersampling, we did not expect to recover the exact times of commitment and progenitor population size. Nevertheless, we wanted to evaluate if these estimates would approach the truth with increased sampling. We thus subsampled the experimental data to lower numbers of cells per terminal state, repeating the process 50 times at varying sample sizes. In parallel, we carried out simulations with the same sample sizes. We then classified inferred fate maps based on their topology and correctness of relative ordering (**Figure 10E**) and found that the fraction of correct topologies and relative order of commitment increased in a similar fashion as more cells were sampled in both simulated and experimental sets. Additionally, commitment times and population sizes of P3 and P4 approached the actual amount with increasing sampling in simulated and experimental sets alike (**Figure 10F,G**). Together, these observations validate our barcoding models used for simulation and indicate that our quantitative fate mapping strategy, ICE-FASE, and Phylotime are robust to the simplifying assumptions made in their models. These results also suggest that developmental differences, ostensibly caused by genetic or environmental factors, can be detected using quantitative fate mapping.

**Figure 10.**
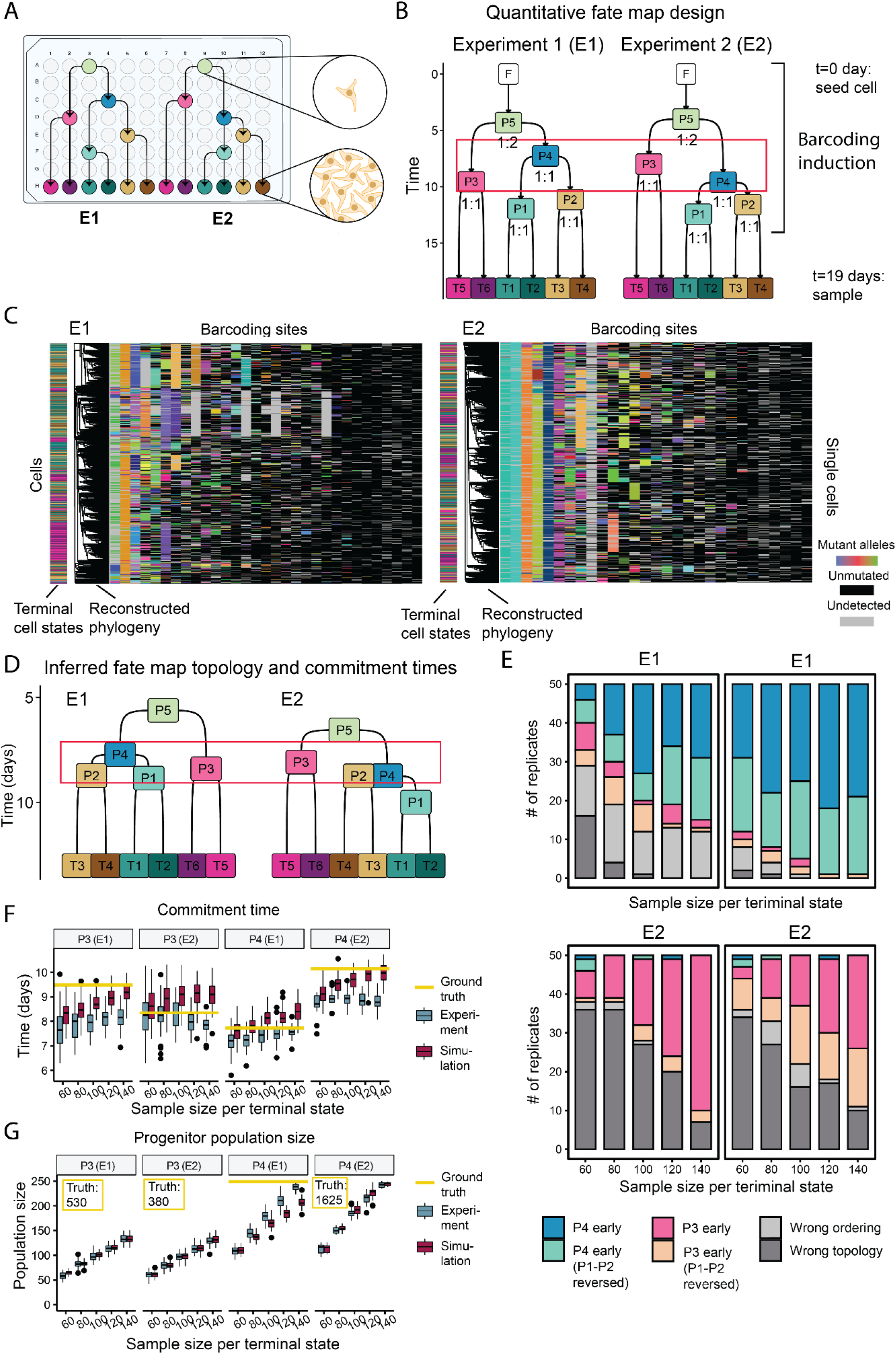
Validation of quantitative fate mapping strategy using an in vitro system. (**A**) Implementation of the split-passage scheme in cultured iPSCs. (**B**) The two quantitative fate maps that were implemented using the passage-split scheme in A. The red box highlights the differences between progenitor state order. (**C**) Barcodes maps from hundreds of sequenced single cells in each experiment are shown (E1 shown on the left, E2 on the right) and used to reconstruct a time-scaled phylogeny which is shown to the left of the map. The state of each cell is marked on the bar to the left of the phylogram and the color code is according to panel B. (**D**) Reconstructed quantitative fate maps from E1 and E2 single-cell barcode results. The red box highlights the differences between progenitor state order. (**E**) Barplot showing the fraction of correctly ordered topologies among a number of subsampled replicates in both simulation and experiment at a number of different sample sizes. **(F)** Boxplot for comparing commitment time estimates and true commitment times for progenitor states 3 and 4 at different sample sizes in simulation and experiment. **(G)** The same comparison for progenitor population size.

## Discussion

In this study, we set out to determine how cell phylogeny—as estimated by lineage barcoding approaches—can be used to understand the dynamics of progenitor states that drive the development of animals. This is a timely problem as advances in genome engineering and sequencing technologies make it possible to estimate cell phylogeny with high throughputs using synthetically-induced or naturally-occurring somatic mutations. However, phylogeny of terminal cells is inherently stochastic and has a complex relationship with the fate of progenitor states. Moreover, retrospective strategies can only sample a fraction of the cells in an organism and are subject to inherent and complex errors in phylogenetic reconstruction.

To address these challenges, we established a robust and practical approach for phylogeny-based analysis of cell fate called quantitative fate mapping. This approach uses terminal cells’ lineage barcode and type information to quantitatively characterize the progenitor field—the collection of progenitor states [20]—that gave rise to them. Our approach first estimates a time-scaled phylogenetic tree based on single-cell barcodes using the Phylotime algorithm. It then combines the phylogenetic tree with the terminal fate of the sampled cells to identify nodes associated with fate decisions and uses their timing to reconstruct an initial map of progenitor states. Finally, it infers the progenitor state of all internal nodes of the phylogenetic tree and uses these assignments to estimate the commitment time, population size, and commitment bias of progenitor states (ICE-FASE algorithm). While other strategies have been described to infer the lineage relationship of terminal cells in specific biological systems, our quantitative fate mapping approach is unique in that it evaluates the dynamics of progenitor states, can be applied to any retrospective strategy, and scales to large and complex fate and lineage landscapes. Moreover, we have demonstrated that this strategy can tolerate the errors that are inherent to phylogenetic reconstruction and used in silico experiments based on realistic barcoding parameters to validate its performance. Importantly, we have used experiments with cultured stem cells to show that our strategy is robust to our simplifying assumptions.

A key finding of our quantitative fate mapping strategy is that time-scaled phylogenies—phylogenetic trees wherein branch length corresponds to actual time—can be used as chronometers of progenitor population dynamics. The few currently available lineage reconstruction strategies that can create time-scaled phylogenetic trees require NP-hard searches in the tree space that become computationally untenable for adequately large sampling depths. To reconstruct time-scaled phylogeny at scale, we developed the Phylotime algorithm which obtains the time since the most recent common ancestor of each pair of cells with maximum likelihood estimation and uses the estimates to reconstruct time-scaled phylogenies. Phylotime performs in polynomial time making it readily applicable to large trees with thousands of terminal branches. Phylotime can be readily adopted for other systems where multiple barcoding sites create recurring mutant alleles in parallel..

This work also establishes a foundation for obtaining meaningful estimates of fate dynamics in a single animal from phylogeny of a small number of cells. Unlike the phylogenetic tree itself, these estimates can be directly combined to create species-level insights or make comparisons between different species. Critically, our results show that only when a progenitor population’ s progeny is adequately sampled can its existence, potency, and quantitative parameters be meaningfully estimated from a single time-scaled phylogeny; estimates in severely undersampled progenitor states are not meaningful and will not be improved by combining multiple animals. To assess if estimates are robust, we defined progenitor population sampling fraction as the fraction of the actual progenitor population whose descendants are observed among terminal cells. We observed that statistically meaningful characterization typically requires sampling fractions larger than 25%. To meet this criterion, the number of terminal descendants that are sequenced should have the same order of magnitude as the progenitor population size (PScov larger than 1). We propose this as a fundamental rule for retrospective lineage analysis approaches. Accordingly, the growing capacity of single-cell sequencing technologies bodes well for quantitative fate mapping since the average progenitor state is less than 1,000 cells during and prior to organogenesis [22,35]. For larger progenitor populations the sampling barrier may be overcome by strategically bottlenecking the number of sampled progenitors. For example, in the neocortex where progenitors develop into known and limited anatomical positions, bottlenecking can be accomplished by sampling a limited number of cortical columns. Alternatively, prospective lineage tracing approaches may be employed to label a subset of a progenitor population, for instance with fluorescence, and only sample their terminal progeny [36].

Sampling strategy is also a critical parameter in experimental design. Our results show that analyzing a fixed number of cells from each terminal state can be more beneficial for characterizing the topology of the fate map especially with more unbalanced topologies, and, to a lesser extent, determining the commitment time of progenitor states. Sampling in accordance with the abundance of each terminal state can be beneficial for determining commitment bias and population size of progenitor states.

Quantitative fate maps obtained in this fashion represent a fate landscape that can capture emergent features of the developmental system that gave rise to the sampled cells. They complement the state manifolds obtained using direct molecular analysis of progenitor populations (e.g., single-cell RNA sequencing) [35,37,38] in multiple ways. Firstly, they provide information about the long-term fate of progenitors. Secondly, they can capture variation between embryos of the same species. Finally, they enable analyzing progenitor populations with respect to a specific subset of their progeny which may be of particular interest, for example due to relevance to a specific genetic or environmental signal.

The limitations of this study are born out of the assumptions of its models. For instance, progenitor states examined here differentiate into only two downstream states. While this assumption does not alter the general conclusions of our study, our strategy can be modified to accommodate other scenarios when necessary. A trifurcation of a progenitor state into three downstream states, for example, can be resolved as two progenitor states with no distance between them on the fate map. Additionally, our models assume parameters of cell division and barcoding mutagenesis tailored to mouse development. These parameters can be adjusted to analyze other organisms, other stages of development, and other developmental systems such as organoids. Importantly, we have not made any assumptions about the mechanisms of differentiation and commitment (e.g., asymmetric versus symmetric cell divisions). As such, the models are agnostic to these parameters and quantitative fate mapping performance may even be enhanced should information about these mechanisms be incorporated. Another limitation of the current model is that it does not consider cell death in progenitor or terminal populations. However, cell death rates would be indirectly reflected in fate map parameters such as population size. Moreover, given some priors about its rate, cell death rates can be incorporated into the model.

In summary, this work provides a framework for high-throughput quantitative fate mapping that can be used both with synthetic lineage barcoding technologies and with naturally occurring somatic mutations. These quantitative fate maps describe progenitor populations that gave rise to sampled cells and characterize their fate dynamics. Robust fate mapping relies on adequate representation of each progenitor population among sampled cells as well as the ability to infer phylogenetic trees with branch lengths that represent time. This work expands the scope of barcoding approaches beyond lineage tracing to quantitative fate mapping which can allow for the characterization of genetic and environmental effects during development.

## Methods

### Definition of quantitative fate map (QFM)

The quantitative fate map describes cell division and fate commitment dynamics at the cell population level. At t = 0, there is one cell at the root (most potent) state. In between fate commitments, at fixed doubling times specific to each progenitor state, cells double to give rise to a new generation of cells at the next time point. Each time there is a fate commitment event, cells transition to one of the two downstream states that are less potent. They do so by first doubling, and have the daughter cells randomly committed to one of the less potent states according to a commitment bias of *p*. Suppose *N* cells commit to downstream states PX and PY. At the commitment time, the *N* cells of the more potent type double once again and [2*Np*] cells become type PX and 2*N* − [2*Np*] cells become type PY. Cells of types PX and PY continue to divide with their respective doubling times until another fate commitment event occurs or the target time of sample collection has been reached.

In summary, parameters of the quantitative fate map include: topology of fate commitment and the following parameters for each progenitor state: doubling time, commitment time, commitment bias. Progenitor population size, defined as the number of progenitor cells at a fate commitment event can be derived from the fate map topology and doubling time. In this work, our interest is in inferring fate map topology, along with commitment time, commitment bias and progenitor population size for each progenitor state.

### Definition of time-scaled phylogeny

A time-scaled phylogeny is defined as a rooted, ultrametric phylogenetic tree where branch lengths are in the unit of time. Nodes in the time-scaled phylogeny represent cell division. The root node represents the most recent common ancestor (MRCA) of all terminal cells. The length of the root edge is the time until the cell division of the root MRCA. Cophenetic distance is defined for each pair of terminal cells, which is the distance between the cells along the phylogenetic tree. The depth of a node in the phylogenetic tree is defined as the distance of a node to the root plus the length of the root edge. The ultrametric property requires that all tips are equidistant from the root, that is, have the same depth. The total time of a time-scaled phylogeny is defined as the depth of its tips.

### Generative model of sampled cell phylogenies based on quantitative fate maps

To generate a phylogeny of a set of sampled cells based on the quantitative fate map, the following steps were employed. First, the number of sampled progenitor cells at each time point were drawn by propagating backward in time, from the terminal cells to the single zygote. At each step, going from some time point *j* to the next time point *j* + 1, suppose *n* cells have divided into 2*n* cells, and *S*_*j*+1_ cells were sampled from the time point *j* + 1. *C*_*j*_ is defined as the number of merges among the sampled progenitor cells at time point *j* + 1, then it can be shown that *C*_*j*_ follows a hypergeometric distribution with the following probability mass function (pmf):

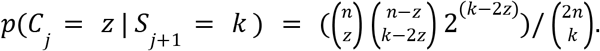

*T*o get the number of sampled progenitor cells in generation *j*, we compute *S*_*j*_ = *S*_*j*+1_ − *C*_*j*_.

Recursively drawing from the above distribution, the progenitor sample sizes at each time point could be derived. At each time point when one cell type commits to two different downstream types, the sample sizes of the first time point of the downstream types were summed, which becomes the sample size of the last time point of the more potent cell type. This process is repeated until the sample size of the zygote is drawn, which is always one.

Next, the exact cell phylogeny is generated based on the progenitor sample sizes. Given that there were *C*_*j*_ = *S*_*j*_ − *S*_*j*+1_ merges at time point *j, C*_*j*_ cells are chosen at random from cells at time point *j* + 1 to be merged. Each merge gives rise to a sampled cell at time *j*. This process allows cells to be recursively merged at each time from terminal population to the zygote. The merges alongside their timing makes up a time-scaled phylogeny.

### Realistic cell division rates during early mouse developement

Before t = 4.2 days, interdivision time was set to 0.6 days. After day 4.2 interdivision time was set to 0.35 day. The number of cells resulting from these division rates were compared to the previously reported numbers in [22]. **Figure S1** compares the number of cells in our simulation based on these doubling times to the estimates of the number of cells in a developing mouse.

### Construction of fate map panel

To generate a series of fate map topologies with varying levels of imbalance, 10,000 random tree topologies were generated using *rtree* function from the “ape” package in R [39], and computed for each category of 16, 32 and 64 terminal states the BSUM index of each tree. Next, the trees were classified into groups with incremental values of BSUM 5. Finally, one tree from each group was randomly selected. The most balanced tree was also appended to each category. The most unbalanced tree for a fixed number of terminal states is pectinate, which is included with this procedure.

The timing of commitment events (bifurcations) given fate map topologies was generated next. To make commitment times comparable across the different categories of fate maps, commitment events were spaced out across a common time window (1.8 - 9.8 days), subject to the constraint of each topology. To do so, a random node order on the topology was first drawn. Specifically, a permutation of the nodes of each daughter was generated at each bifurcation, and subsequently combined [40]. The ordering ranked each commitment event from earliest to latest. Next, the intervals between consecutive commitment events were taken as the parameters, and we minimized the values of these parameters, subject to the constraints of fate map topology and allowing for at least one cell doubling in between two consecutive commitments, using linear programming optimization. The remaining duration of the interval in addition to the minimum length was then distributed into each interval according to a symmetric Dirichlet distribution with *α* = *5*. Finally, the commitment biases in each fate map were drawn from a Beta distribution with *α* = *β* = *5*.

### Fate map topology reconstruction with FASE

To determine if a node is a FAte SEparation (FASE), all unique terminal types of the progeny of each node were listed. The node was classified as a FASE (for at least some pair of terminal types) if any of its daughter nodes had observed fates that are less potent. We identified FASE nodes across all nodes in the phylogenetic tree. Next, for each FASE, all pairs of terminal states that a FASE separated were listed. Now for a pair of terminal fates, the mean depth of the FASEs that separated the terminal fates were computed, referred to as the FASE time. If no FASE existed for a pair of terminal states, the FASE time was taken to be zero. Finally, the distance between a pair of terminal states was equal to two times the difference of total time and the FASE time. To get the topology from the full distance matrix, the *upgma* function from the “phangorn” package [41] was applied, which is a wrapper of *hclust* in base R.

### Node state assignment for time-scaled phylogeny

Each bifurcation of the fate map corresponds to the commitment event of a progenitor state. One characteristic of the progenitor state is its potency: the set of terminal states it can lead to. Each node in the time-scaled phylogeny also had an observed potency determined by the set of states its progeny covers. The progenitor state of a node of the phylogeny was assigned greedily based on potency: the node was assigned a state that had the same potency as itself. If no such state exists in the fate map, then it was assigned a least potent state in the fate map that was more potent than the node.

### Commitment time inference with ICE

To infer commitment time of a progenitor state, inferred commitment events (ICEs) were identified. A node in the time-scaled phylogeny was considered an ICE if both daughters had a different assigned state than itself. Each ICE was associated with a progenitor state. The depths of all ICEs associated with a state defined the ICE times. The mean of ICE times was used as an estimate for commitment time. In cases where inferred commitment times of the downstream state is earlier than that of the upstream state, the commitment time of the downstream state was set to that of the upstream state.

### Progenitor population size and commitment ratio inference

To identify the population present at the commitment time of a progenitor state, a set of branches in the time-scaled phylogeny needed to be identified. First, a set of extended states was defined. The extended states included the state itself, its upstream states up to root and its downstream states down to the terminal states. Next, a state path was constructed for each branch that spanned the commitment time of the progenitor state: the state path included a number of connected states on the fate map topology, which started at the state of the incoming node and ended at the outgoing node of the edge. A branch was considered associated with the progenitor state if its state path overlapped with any of the extended states. To estimate the progenitor population size, the collection of incoming nodes of the associated branches were counted.

Next, to quantify the bias of a progenitor state’ s commitment, each associated branch was further classified into one of the three categories based on if it was committed to one or the other immediate downstream states, or if it was uncommitted. The classification was made based on the state path; if the state path covered one of the two immediate downstream states, it was classified accordingly. Otherwise, it was classified as uncommitted. To estimate the commitment bias, the ratio of associated branches committing to one side versus the other was used.

### Estimation of mutagenesis parameters in MARC1 mice

To get posterior estimates of mutation rates of MARC1 hgRNAs (i.e., λ, rate of the Poisson process), a grid search was conducted to match empirical distributions of mutated fractions among simulated and observed data across a number of embryonic time points. Previously reported hgRNAs formed three classes: the ‘ slow’ class generated mutations on the order of 0.001 mutations/day, ‘ mid’ class generated ∼0.1 mutations/days, and fast class generated ∼1.0 mutations/day during early mouse development. The ‘ slow’ and ‘ fast’ estimates expectedly had large uncertainties as most observed fractions are close to 0 or 100 percent mutated. Alternatively, a naive estimate of mutation rate can also be used. If mutated fractions *F_i_* were observed at time *T*_*i*_ in animal *i* for*i* = 1,…, *N*, then 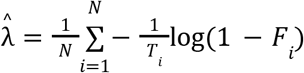 is a naive estimate.

For mutant alleles of a barcoding site, estimating the probabilities of individual repair outcomes created by Cas9 DNA break-repair (mutant emergence probabilities) was challenging. Normally, the fraction of cells carrying a particular mutant allele among all cells with a mutated allele (within-animal estimates) is a good estimator of the allele emergence probabilities. However, when cells divide and mutate starting from a small field size, these fractions are largely affected by the time of the mutagenesis events, as early events result in larger clones carrying the mutation. On the other hand, when hgRNA genotypes are observed for multiple animals, the fraction of animals that carry a particular genotype, once normalized, and when the probability is small, can be good estimates to the mutation probabilities (across-animal estimates) (**Figure S3B**,**C**). In this case, the estimation accuracy depends on the number of animals analyzed. From the MARC1 time course data, the within-animal estimates were calculated for each animal and averaged, and the across-animal estimates were calculated based on 173 embryos from 2 mouse lines. To get a more complete profile of possible mutant alleles and their occurrence probabilities for each hgRNA, we adapted the inDelphi machine learning algorithm to predict CRISPR-Cas9 mutation results [30] for hgRNAs. We observed that the inDelphi-predicted probabilities agreed well with the across-animal estimates from MARC1, but poorly with within-animal estimates (**Figure S3B**). Further, the fact that the majority of the low probability mutations were not observed in any mouse suggests that the limited number of mutagenesis events during mouse development does not sufficiently cover the majority of the mutational profiles. These conclusions were further validated by simulating multiple animal lineage barcode data based on inDelphi-predicted mutational profiles and comparing the within- and across-animal estimates of the true parameters (**Figure S3C**).

### InDelphi predictions of hgRNA allele emergence probabilities

The emergence probabilities of hgRNA mutant alleles were computed by inDelphi. inDelphi is a machine learning algorithm to predict heterogeneous insertions and deletions resulting from CRISPR/Cas9 double-strand break. [30] In this study, inDelphi model trained with the mouse embryonic stem cell mutation dataset was used to predict the probabilities of hgRNA mutants from MARC1 mice. The original 64 hgRNA sequences in MARC1 mice were subjected. Since Cas9 nuclease cuts 3 bp upstream of the Protospacer Adjacent Motif (PAM, NGG sequence)^1^, the possible mutations from the cut site at −3 bp from the PAM sequence were computed. To take into account the repeated targeting of hgRNAs, inDelphi is first applied to predict a set of first-round mutations. Subsequently, the resulting first-round mutations were used as inputs to the next round of inDelphi predictions. Notably, only mutant sequences with >16 bp protospacer and PAM are subject to the second round analysis as gRNA without >16 bp spacer sequence loses its activity [42]. Here, the probabilities of the next generation mutants were computed by multiplying the score of the mutant by the score of the parental mutant. Repetitive application of inDelphi produces exponentially growing numbers of potential mutant alleles. Therefore, the analysis was limited to three cycles, resulting in first to third generations of mutants. Since the different generations can have the same mutant sequences, the scores corresponding to the same sequences were summed. Finally, probabilities of all mutant alleles were normalized to have a sum of one.

### Simulation of lineage barcodes from time-scaled phylogeny

To simulate lineage barcodes mimicking barcoding in MARC1 mice, 100 hgRNAs that were of the ‘ mid’ or ‘ fast’ fast category were randomly sampled from the MARC1 pool of hgRNAs for each simulation (**Supplementary Data 4**). The corresponding estimated mutation rates and inDelphi predicted allele emergence probabilities were used as input to the mutagenesis model.

### Phylotime for reconstructing time-scaled phylogeny from lineage barcodes

Our approach to reconstructing time-scaled phylogenetic trees for thousands of cells is based on a maximum likelihood estimation of pairwise temporal distances between cells. Given a pair of terminal cells (**Figure S8**), we estimated the branch length parameter 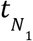, which is the time since the most recent common ancestor (MRCA) of the two cells. For a barcoding site*i* with a mutation rate of λ_*i*_, and probabilities *a*_*i*_ = *(a*_*1*_, *a*_*2*_,…, *a*_*J*_ )_*i*_ of mutating into alleles *(A*_*1*_, *A*_*2*_, …, *A*_*J*_)_*i*_, the likelihood of observing a given allele in a single barcoding site in two cells (*c*_1_, *c*_2_) is the sum of two terms:

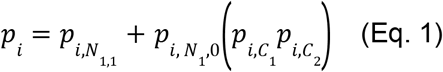

The first term, 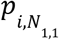, is the probability that a mutation has occurred before the MRCA, leading to identical alleles in both cells. The second term is the probability of observing the allele in each terminal cell respectively (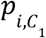and 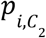) conditional on no mutation occurring before the MRCA 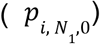. Because barcoding sites in our model were independent, the likelihood for the set of alleles observed in all barcoding sites was then the product of their individual likelihoods:

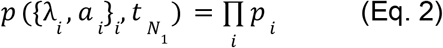

*T*o get estimates of pairwise distance, or equivalently, time until MRCA between two cells 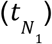 we first plugged in estimates of mutation rates and allele emergence probabilities. Here, naive estimates of mutagenesis parameters were plugged in. The estimates of mutation rates were obtained by 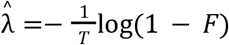, where *F* is the mutated fraction and *T* is the total time from the start of the experiment to the sample collection. Mutant allele emergence probabilities were set according to a uniform prior, that is *a*_*j*_ = 1*/J* for *j* = 1,…, *J*. Alternatively, better estimates such as the across-animal estimates or inDelphi predictions may be used and are expected to improve Phylotime performance. To get the optimal value of 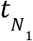, the following score equation was solved using the Newton-Raphson method:

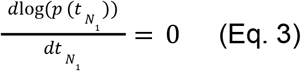

*T*he distance between the two cells is 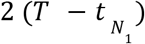. Once all pairwise distances were computed

*w*ith the above method, we applied UPGMA hierarchical clustering [32] to derive a phylogenetic tree wherein branch lengths represent actual time. We called this approach Phylotime.

### Generation of an inducible Cas9 barcoded stem cell line

#### Stem Cell Culture

Human induced pluripotent stem cells (iPSCs) from the EP1 line [43] were cultured in mTeSR1 (STEMCELL Technologies) on plates coated with Matrigel Growth Factor Reduced Basement Membrane Matrix (Corning). Cells were maintained at 37°C and 10% CO_2_/5% O_2_ conditions with daily media changes. When up to 80% confluent, cells were passaged by dissociation with Accutase (Sigma Aldrich) and seeded in mTeSR1 media supplemented with 5 μM blebbistatin (Sigma Aldrich).

#### Knock-in of an Inducible Cas9 Cassette

EP1 iPSCs were modified to express Cas9 protein under doxycycline induction. CRISPR/Cas9 was used to target and insert both a reverse tetracycline-controlled transactivator (rtTA) construct and a tetracycline-dependent Cas9 construct into each of the two copies of the AAVS1 safe harbor locus. The following plasmids were used to generate the cell line:

*Cas9/AAVS1 gRNA*-modified pSpCas9(BB)-2A-Puro (PX459) V2.0 (Addgene Plasmid #62988 [44] with T2A replaced with P2A) with gRNA sequence caccGGGGCCACTAGGGACAGGAT

*Cas9 donor*-modified Puro-Cas9 donor (Addgene Plasmid #58409 [45] with puromycin resistance replaced with blasticidin resistance)

*AAVS1 donor*-AAVS1-Neo-M2rtTA (Addgene Plasmid #60843 [46])

Cells were grown to 80% confluency and then dissociated with Accutase for 13 minutes to generate a single cell suspension. 50,000 cells were resuspended in mTeSR1 media with 5 μM blebbistatin and seeded into one well of a 24-well plate coated in Matrigel. The following day, 350 ng of Cas9/AAVS1 gRNA plasmid and 500 ng each of Cas9 donor and AAVS1 donor plasmids were combined and added to 48 ul Opti-MEM (Gibco). 2 ul of Lipofectamine Stem Transfection Reagent (Thermo Fisher) were added to the transfection mix, which was then vortexed and incubated for ten minutes at room temperature. The entire transfection mix was added to one well of cells. Media was replaced the following day. 40 hours after transfection, cells were transiently selected for 24 hours with 0.95 ug/mL puromycin, 5 ug/mL blasticidin, and 200 ug/mL G418 sulfate. Surviving cells were cultured to at least 30% confluency, and then dissociated to a single cell suspension for clonal expansion. 500-1000 cells were seeded in one well of a 6-well plate and cultured for 7-10 days before clonal colonies were picked and screened for the intended insertions. PCR was performed with the following primers to confirm knock-in of the Cas9 and rTTA constructs:

*Cas9_F*: CACCTTGTACTCGTCGGTGA

*rtTA_F*: GCTGATTATGATCCTGCAAGC

*AAVS1_RHA_R*: GGAACGGGGCTCAGTCTGA

Positive colonies were cultured and clonally expanded once more, with a second round of colony picking and PCR screening to ensure clonality of the final cell line.

#### Lentiviral Infection with an Array of hgRNAs

50,000 cells from the doxycycline-inducible Cas9 line were seeded into one well of a 24-well plate. The following day, 300 ng of Super PiggyBac Transposase (SBI System Biosciences), 700 ng of PiggyBac-hgRNA L21 library ([9]), and 50 ng of PiggyBac-hgRNA L21 + puro library were combined and added to 48 ul Opti-MEM (Gibco). 2 ul of Lipofectamine Stem Transfection Reagent (Thermo Fisher) were added to the transfection mix, which was then vortexed and incubated for ten minutes at room temperature. The entire transfection mix was added to one well of cells. edia was replaced the following day. Transfected cells were selected with 0.95 ug/mL puromycin for one week.

Selected cells were dissociated to single cell suspensions, and 500-1000 cells were seeded in one well of a 6-well plate and cultured for 7-10 days before clonal colonies were picked. Colonies were screened for hgRNA insertions using qPCR. Genomic DNA from each colony was amplified with the following pairs of primers:

*Sox11_F*: TGATGTTCGACCTGAGCTTG

*Sox11_R*: TAGTCGGGGAACTCGAAGTG

*hgRNA_F*: ATGGACTATCATATGCTTACCGT

*hgRNA_R*: TTCAAGTTGATAACGGACTAGC

For each colony, the cycle threshold value for hgRNA amplification was subtracted from that of Sox11 amplification. Colonies with the largest cycle threshold value difference, indicating the highest number of hgRNA insertions, were cultured and clonally expanded once more. Colonies were picked one additional time to ensure clonality of the final cell line.

#### Determining cell line hgRNA array identity and function

Cell line hgRNA identities and functions were determined by performing a doxycycline time course experiment. Genomic DNA was extracted from cells after 0, 4, 8 and 11 days of doxycycline treatment and Cas9 induction. hgRNA sequencing libraries were prepared, sequenced, and analyzed as per the published pipeline [28]. The percent of reads for each hgRNA identifier sequence that were mutated was calculated at each time point (**Figure S4**), determining the relative activity for every hgRNA.

### In vitro quantitative fate map experiments

Single cells from the Cas9-hgRNA iPSC cell line were FACS sorted into a 96-well plate coated with Matrigel and containing mTeSR plus medium supplemented with 5 μM blebbistatin, 10% CloneR (StemCell Technologies), and 1 μM Pifithrin-α hydrobromide (Tocris), 1X Antibiotic-Antimycotic (Gibco), and 0.2 ug/uL doxycycline. Supplemented media was replaced every other day. Wells were assigned to follow specific quantitative fate maps, and were passaged at times, sizes, and ratios as determined by the fate maps. To passage small numbers of cells, media was aspirated from wells and 30-100 ul of Accutase (depending on the size of the well) was added and incubated for 8 minutes. The Accutase and cell suspension was directly added to supplemented media in the wells the cells were passaged to. Passaged cells were incubated for 2 hours to allow cells to attach to the Matrigel coating, after which a 50% media exchange was performed to decrease the amount of Accutase remaining with the cells. At the end of the experiments when the cells were passaged into their terminal wells, doxycycline treatment was ended so that barcode editing would discontinue.

### Determining progenitor population size from in vitro experiment

Brightfield images were taken of cells at P5, P4, and P3 for E1 and E2 before passaging (**Figure S6**) To estimate the cell numbers at each state, images were analyzed in ImageJ. Outlines were drawn for 5 different cells, and the average area noted. The total area of the colony or colonies was then measured, and divided by the average cell area to estimate the total number of cells present in each image.

### Sequencing single cell lineage barcodes

Cells were dissociated into single cell suspensions and diluted in PBS pH 7.4. Single cells were FACS sorted into 1 ul each of QuickExtract DNA Extraction Buffer (Lucigen). DNA was extracted by incubating cells for 10 minutes at 65°C followed by 5 minutes at 98°C. Single cell barcode sequencing libraries were generated using serial PCR reactions as per the published protocol ([28]) with each cell treated as an individual sample. Libraries were sequenced on a MiSeq System (Illumina).

### Data processing for in vitro experiment data

Identifier and spacer sequence pairs were extracted for each sample using step one of the published pipeline. [28] For cells that were sequenced more than once, “identifier+spacer” pair counts were first merged for each cell. The merged data were then provided as input to the remainder of the pipelines for sequencing error correction and filtering.

A total of 32 hgRNAs were identified, each observed in more than 921 cells. Sequencing errors among identifier sequences were first corrected. For each identifier sequence that was not one of the 32 true hgRNAs, if the identifier sequence was within a hamming distance of 1 to any true identifiers, its spacer counts were merged with that of the true hgRNA’ s. After the correction, no other identifiers other than the 32 true hgRNAs were observed in more than 3 cells.

Spacer sequencing errors were corrected next. First, the error reads within each cell and hgRNA combination were corrected. In one of our sequencing runs, one cycle of sequencing returned ‘ N’ for all spacer sequences. These errors were computationally corrected: if there existed another spacer sequence for the same identifier and cell that was exactly the same except for the ‘ N’ base pair, the count of the error spacer with the ‘ N’ base was merged with the other spacer. Next, the spacer sequencing errors across different cells were corrected. Again, the error involving ‘ N’ base pairs were further corrected across the cells using the same criteria as the within cell correction.

Each allele was labeled as unmutated or mutated by comparing the spacer sequences to that of the reference sequencing result, that is, if a spacer sequence was observed in the parent for the same identifier, it was labeled as unmutated. One identifier “GCCAAAAGCT” did not amplify in the parent data, and the sequence “GAAACACCGGTGGTCGCCGTGGAGAGTGGTGGGGTTAGAGCTAGAAATAG” was identified as the unmutated spacer based on alignments of its different observed alleles.

Noisy reads were further filtered for cell+hgRNA combinations that had more than one spacer observed. If the most abundant spacer was at least four times more abundant than all the other spacer reads observed, only the most abundant spacer was kept. All the spacer counts with fewer than five total reads were also excluded.

After processing, 1197 cell+hgRNA combinations out of the total 54,012 (2.2%) still had more than one spacer observed. If more than two spacers were observed for more than two hgRNAs in a single cell, the cell was likely a doublet. 33 such cells were identified and filtered. Each cell had a median 25 out of 32 hgRNAs detected. Each hgRNA was detected in a median of 83% of all cells (**Figure S7**). Before reconstruction, non-informative cells and hgRNAs were filtered out. Any hgRNA with a diversity of one, that is, all cells in which an hgRNA was observed had the same allele, was considered non-informative and was excluded. An allele was considered informative if it was mutated and observed in more than one cell in each group, and cells with less than three informative alleles were filtered out. In all, 970 out of 1051 cells for 31 hgRNAs passed the filters for E1 and 943 out of 1032 cells for 29 hgRNAs passed filters for E2.

### Imputation of missing hgRNA alleles with xgboost

As an intermediate step, the missing alleles were imputed before ICE-FASE could be applied to the in vitro data. First, the missing percentages of all hgRNAs were computed, and hgRNAs were imputed one by one from the least amount of missing to the most amount of missing. To impute missing alleles for a single hgRNA, an ‘ xgboost’ model with multinomial softmax objective was trained using all the observed cells as training [34]. The model classified each missing cell as one of the observed alleles. The design matrix was constructed where each column is a mutant allele observed in one of the other hgRNAs. The parameters ‘ max_depth=4’ and ‘ nrounds = 20’ were used for imputing the in vitro data.

### Ground truth fate map of in vitro experiment

To conduct simulations that best resembled the in vitro experiment, the effective cell division rates during the experiment were first determined. First, the division rate of P5 was chosen so that the population size at the first split agreed with what was observed. Next, we assume that cells were split in proportion to the volume of the suspension at the split. The division rate of P3 and P4 were set so that their respective population sizes at the split agreed with what was observed. The division rate for P1, P2 and all the terminal cells were set to once every 20 hours. The exact division rate chosen and the population size at each stage are detailed in **Figure S5**. Notice that the commitment happens one cell division prior to the well split, so the ground truth commitment time is one cell division earlier than the spit time, and the progenitor field size is half of what is observed in the well at the split.

## Acknowledgements

The authors would like to acknowledge Kian Kalhor, Dr. Yuxin Zhu, Dr. Justus Kebschull, and Dr. Loyal Goff for comments on the manuscript. This work was supported by grants from the Simons Foundation (SFARI 606178) and the National Institutes of Health (NIH) (U01HL156056, R01HG009518, R01HG010889, P30EY001765, F31EY030769). Parts of this work were carried out at the Advanced Research Computing at Hopkins (ARCH) core facility, which is supported by the National Science Foundation (NSF) grant number OAC 1920103. Related research in the Kalhor Laboratory is supported by a Packard Fellowship for Science and Engineering. Related research in the Zack Laboratory is generously supported by Research to Prevent Blindness and the Guerrieri Family Foundation.

## Author Contributions

W.F. and R.K. conceived the project; W.F. established statistical frameworks, carried out computational analyses, and designed experiments; C.M.B. created and characterized barcoding cell lines and carried out in vitro experiments with assistance from W.F. and K.L.; A.S. helped create the maximum likelihood framework for Phylotime; S.A. created the inDelphi-based allele predictions; R.K. and W.F. interpreted the data and wrote the manuscript with assistance from C.M.B. and A.S.; all authors commented on the manuscript; D.J.Z, H.J., and R.K. supervised the project.

## Additional Information

Codes and data are available from https://github.com/Kalhor-Lab/QFM/ or upon request.

