## Supplementary Figures and Tables for "Quantitative fate mapping: Reconstructing progenitor field dynamics via retrospective lineage barcoding"

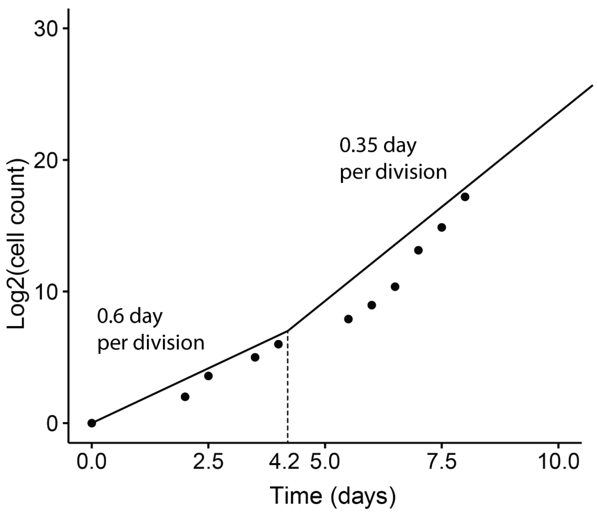


**Figure S1**. Comparison between the total number of cells in simulation and the reported number of cells in early mouse embryogenesis. Dots represent the number of cells from Kojima et al., 2014. The line represents the number of cells in the simulation. The two cell division rates used for simulation are shown.

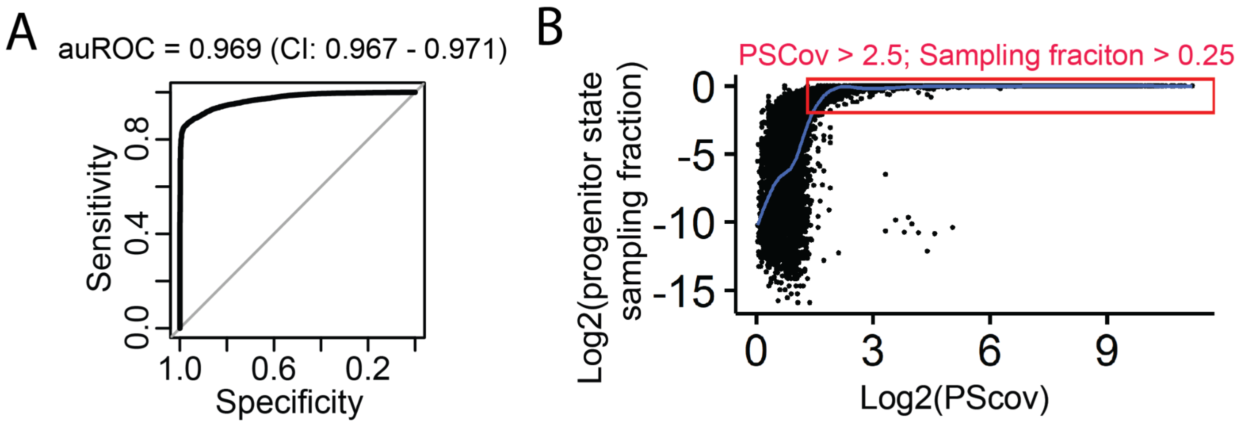


**Figure S2**. Progenitor state coverage statistics (PScov) reveal robustness of obtained quantitative fate map parameters. (**A**) Area Under the Receiver Operating Characteristics (auROC) curve for progenitor state coverage shows a high sensitivity and specificity in detecting adequately sampled progenitor states. CI: confidence interval. (**B**) Scatter plot of progenitor state sampling fraction as a function of its PScov among 508 simulated experiments. Progenitor states with sampling fraction better than 0.25, which produced robust estimates of progenitor states, tend to have coverage above 2.5, as highlighted by the red box. Trendline (LOESS) is shown in blue.

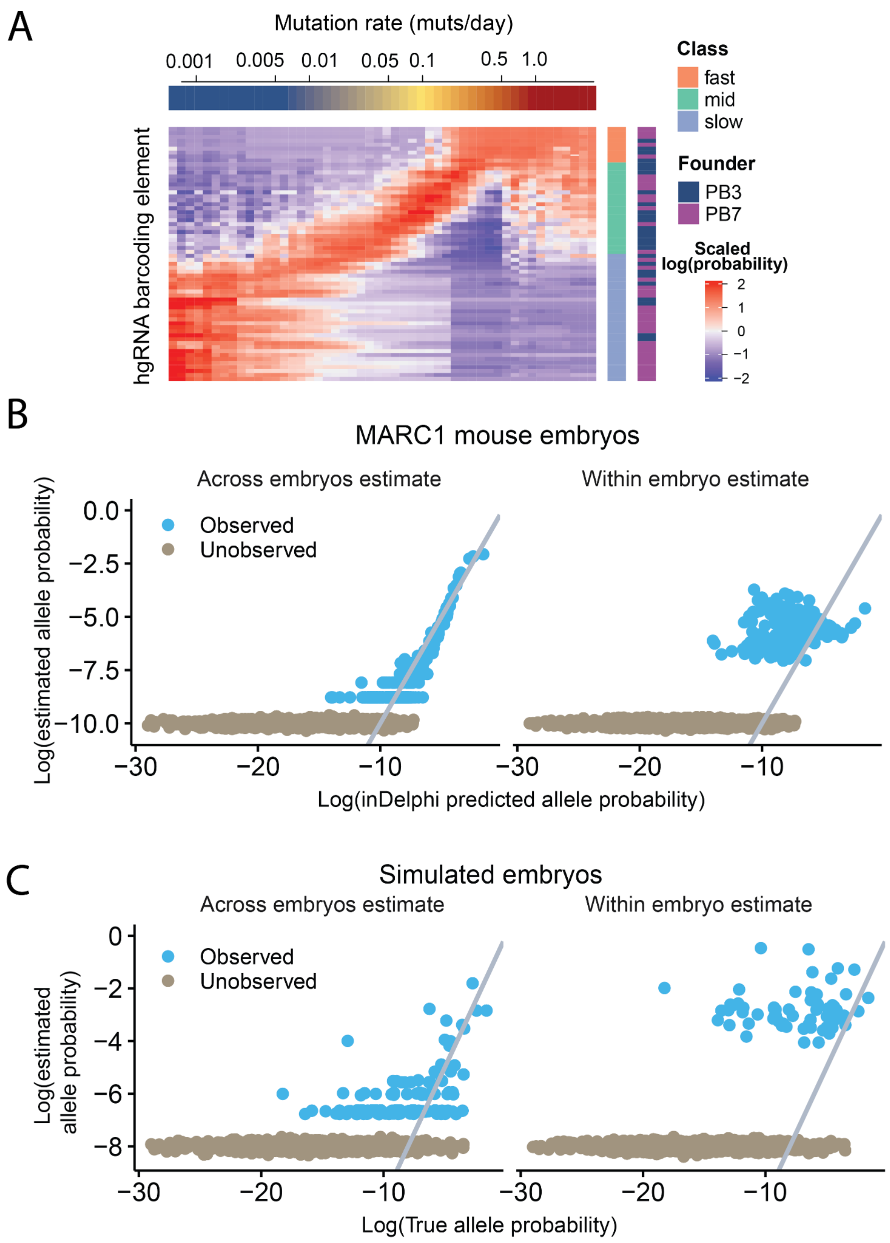


**Figure S3**. Agreement between inDelphi-based allele predictions and those observed in simulation and mouse experiments. (**A**) Posterior probabilities of mutation rates for all MARC1 hgRNAs which originate from either the PB3 founder mouse or the PB7 founder mouse (Leeper et al., 2021) as shown in either dark navy or purple on the color bar to the right. Also shown is the initial characterization label of the hgRNA as either fast, mid, or slow (Leeper et al., 2021). (**B**) Comparison of mutant allele probability estimates of our modified inDelphi algorithm to those estimated from MARC1 mouse embryos by taking either abundance within the embryo when the allele is present (right) or fraction of embryos that present the allele (left). Blue dots represent predicted alleles that were observed in MARC1 embryos, gray ones were predicted by modified inDelphi but not observed in embryos. Alleles observed in embryos but not predicted by inDelphi, which were a very small fraction, are not shown. Gray line has intercept 0, and slope 1. (**C**) Comparison of true mutant allele probability as estimated (x-axis) with their average observed fraction in simulated mouse embryos (right) and fraction of simulated embryos that showed the allele (left). Blue dots represent alleles that were observed in simulated embryos, gray ones were possible in simulation by not observed in simulated embryos. Simulations were conducted 100 times for 9 time points. Gray line has intercept 0, and slope 1. Panels B and C show that the estimator based on allele occurrences across multiple samples outperforms average allele fraction in both simulation and MARC1 mouse data.

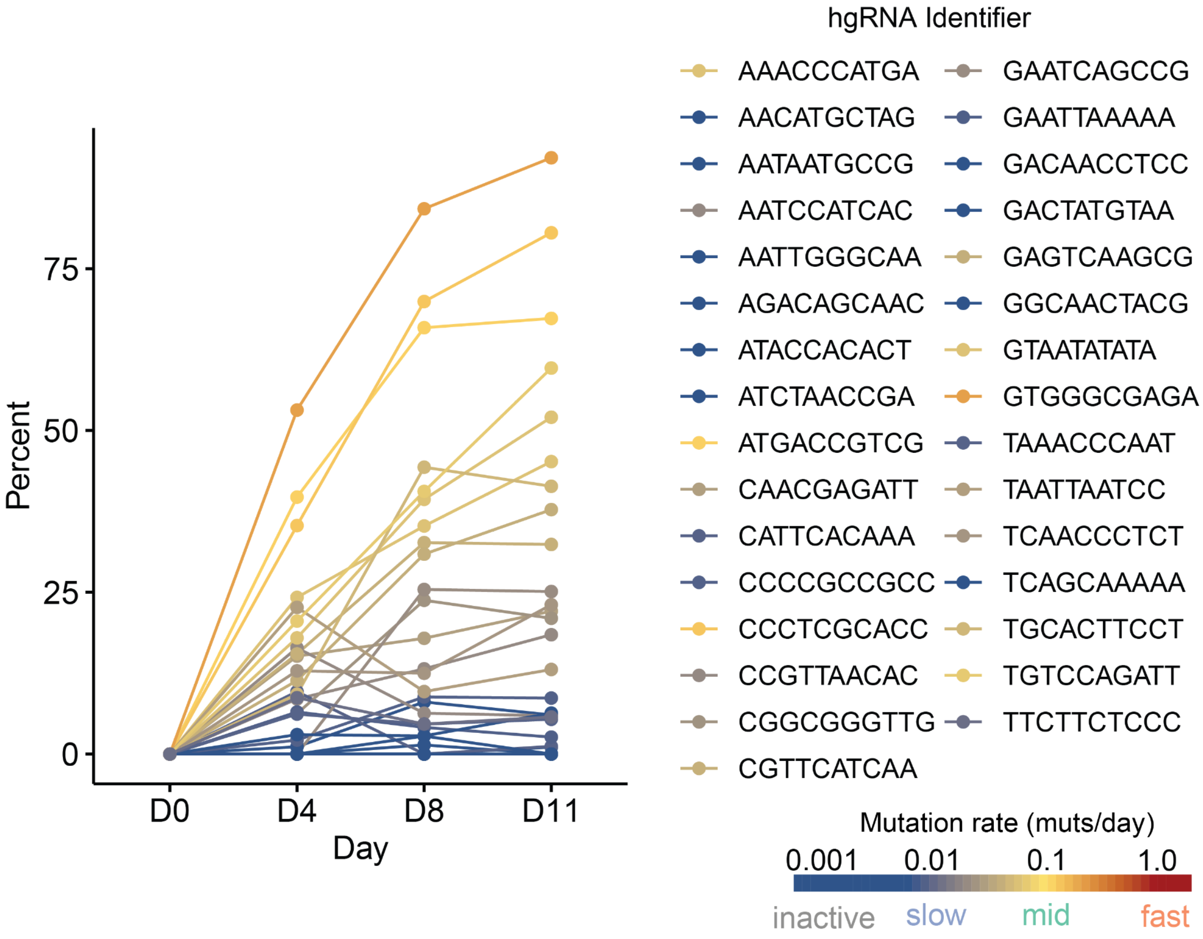


**Figure S4**. Mutated fraction of hgRNAs over time in the iPSC line. Color shows the estimated mutation rate for each hgRNA according to the key on bottom right. The list on top right denotes the color of each hgRNA on the plot.

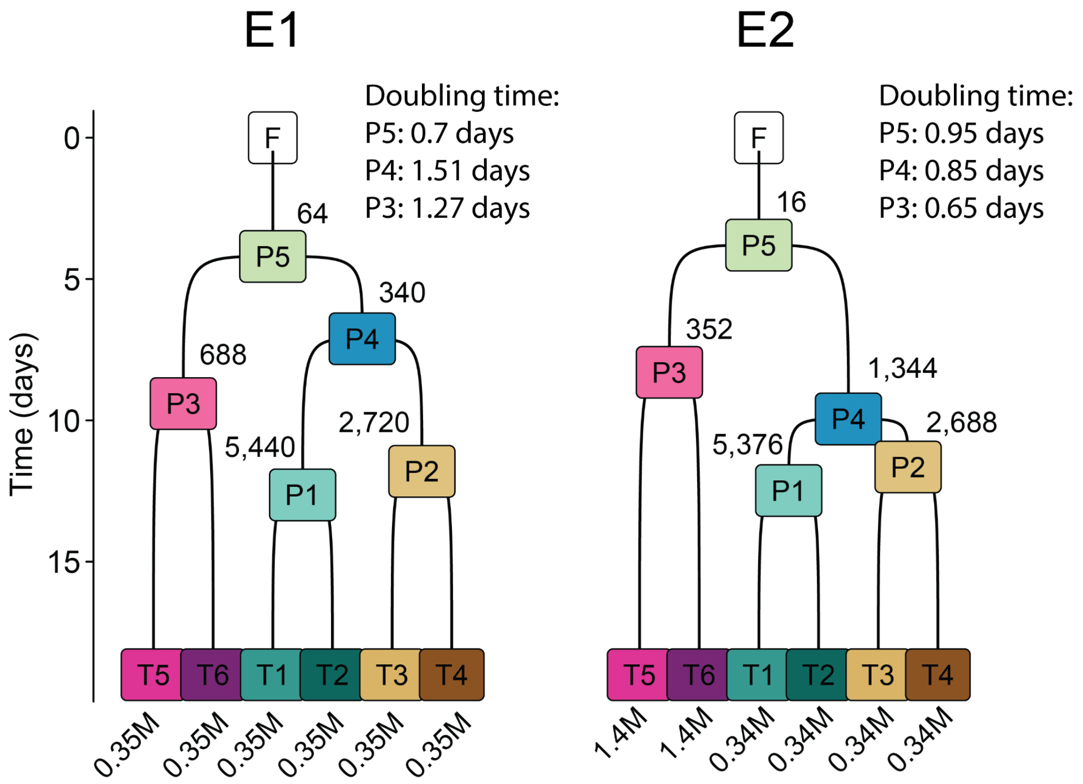


**Figure S5**. Cell division rates and progenitor population sizes in the ground truth fate maps for in vitro experiments. Numbers on the top right corner of each progenitor state show its total population size estimated from bright field images. Numbers on the bottom of each terminal state show its estimated population size from bright field images. The estimated doubling times, which represent cell division rates, estimated from population size at different times, are shown on the top right of each map.

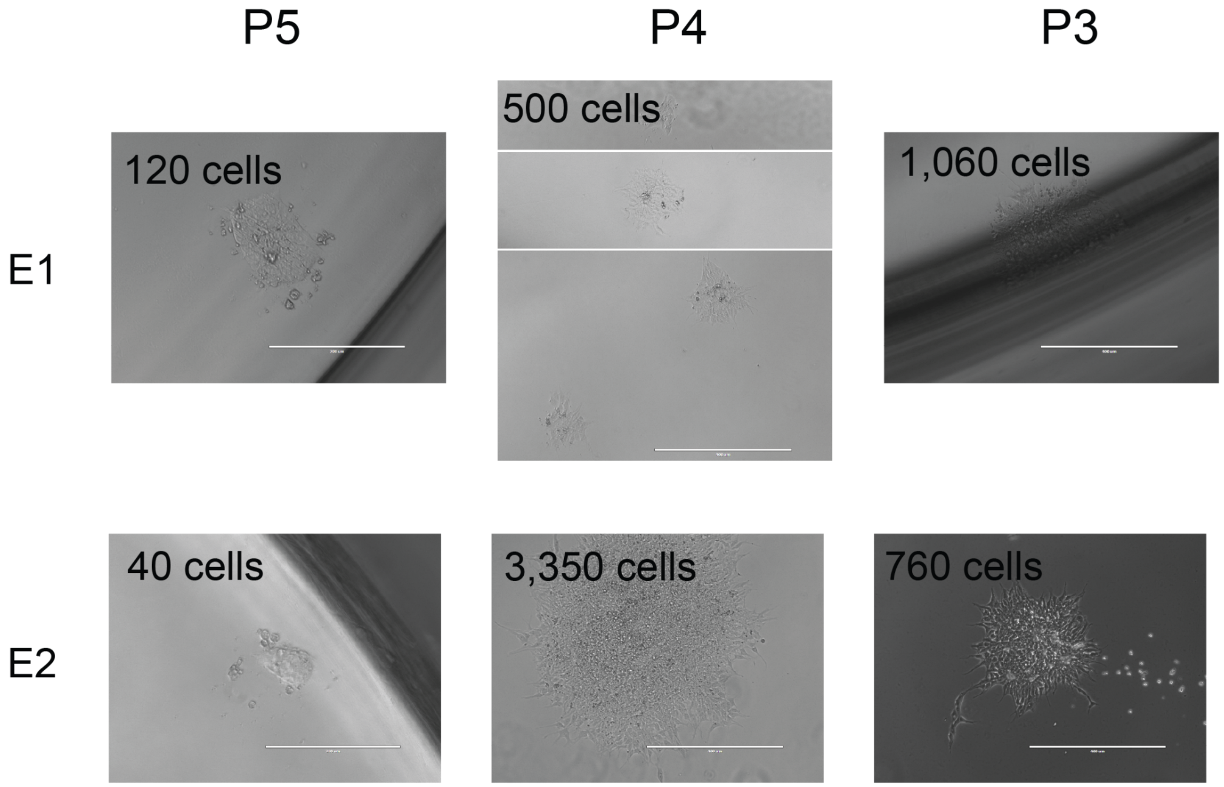


**Figure S6**. Bright field images showing the P3, P4, and P5 progenitor population size estimates (columns) for E1 and E2 experiments (rows). Scale bars for P5 images are 200 microns. Scale bars for P4 and P3 images are 400 microns.

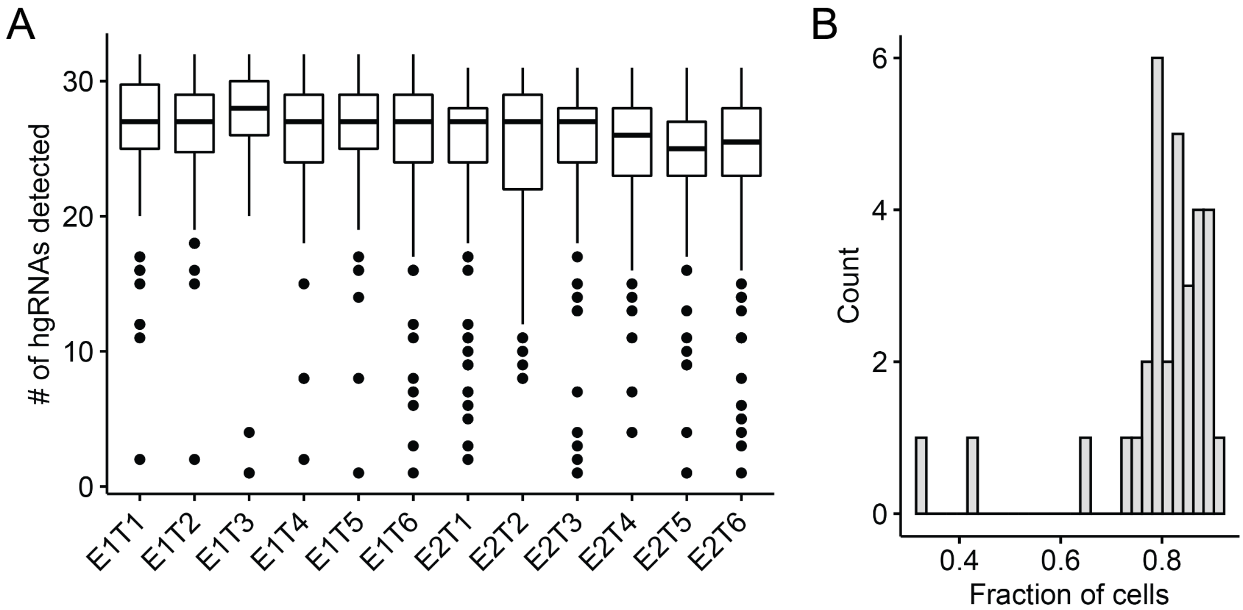


**Figure S7**. Amount of undetected hgRNA alleles. (a) Boxplot showing number of hgRNAs detected (out of the total 32) in each terminal well of each experiment. (b) Histogram of the fraction of cells in which each hgRNA was detected.

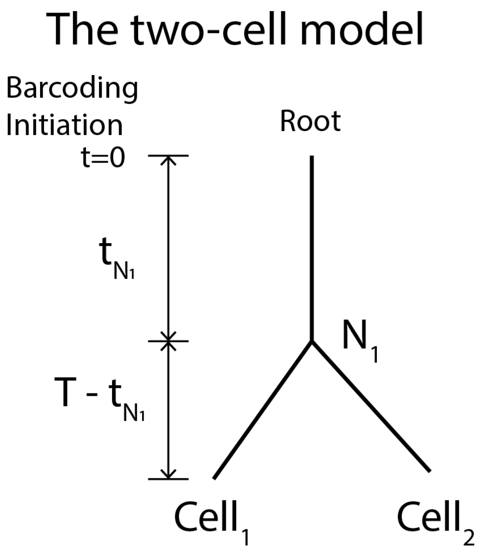


**Figure S8**. Diagram showing a two-cell phylogram illustrating Phylotime likelihood computation. $N_1$ is the most recent common ancestor (MRCA) of terminal $Cell_1$ and $Cell_2$.

**Table S1**. hgRNA identifier and spacer sequences amplified from the iPSC line.

| hgRNA Identifier Sequence | hgRNA Spacer Sequence |
|---|---|
| AAACCCATGA | GGTTTAGTATGGAGGGAAGTG |
| AACATGCTAG | GGTTTTCGATCATAGGTCGTG |
| AATAATGCCG | GGTGCCCTTTATGGGACCGTG |
| AATCCATCAC | GGTGACAAACGTATGTCAGTG |
| AATTGGGCAA | GGTTGTGGGGAACTCGGTGTG |
| AGACAGCAAC | GGTGTTCTTTGCTCGAGGGTG |
| ATACCACACT | GGTTGTAAGCGATGGTTTGTG |
| ATCTAACCGA | GGTGTGCGGAGATTATGCGTG |
| ATGACCGTCG | GGTCCTAGAAGCTGAGTTGTG |
| CAACGAGATT | GGTGCTAGCGGTTTCGAAGTG |
| CATTCACAAA | GGTGGCAATTCGATCTGAGTG |
| CCCCGCCGCC | GGTAGAATGGATCCACGGGTG |
| CCCTCGCACC | GGTTCAACCGGCGTCTTTGTG |
| CCGTTAACAC | GGTTGGCTTTACTCCTTTGTG |
| CGGCGGGTTG | GGTCGTCGTTCGTAGGGCGTG |
| CGTTCATCAA | GGTGAGAACAGAACGTTTTGG |
| GAATCAGCCG | GGTTTTAGTAAATGGTGAGTG |
| GAATTAAAAA | GGTGCTAATAGTTAGCTCGTG |
| GACAACCTCC | GGTAATCTAAAGATCCCCGTG |
| GACTATGTAA | GGTACTAGTTACTTACGGGTG |
| GAGTCAAGCG | GGTTATCGTTACGGATTTGTG |
| GCCAAAAGCT | GGTGGTCGCCGTGGAGAGGTG |
| GGCAACTACG | GGTGGCAGCCAGCCACTTGTG |
| GTAATATATA | GGTGAGCATGATAACGTCGTG |
| GTGGGCGAGA | GGTGAGATGCCTCAAGTGGTG |
| TAAACCCAAT | GGTTTACTTAGTTAACTAGTG |
| TAATTAATCC | GGTTGAGATAATCAAAAAGTG |
| TCAACCCTCT | GGTGACGCGAGGACGGTGGTG |
| TCAGCAAAAA | GGTG |
| TGCACTTCCT | GGTTCCCGTTAGCTCGTAGTG |
| TGTCCAGATT | GGTCTTCCTGGAATCTACGTG |
| TTCTTCTCCC | GGTCGCGAAAATGTGCCGGTG |

### List of supplementary data

Supplementary Data 1. All quantitative fate maps.

Supplementary Data 2. Mutation rate estimates for hgRNAs in MARC1 mice and iPSC line.

Supplementary Data 3. inDelphi predicted mutant allele probabilities for hgRNAs in MARC1 mice and iPSC line.

Supplementary Data 4. Simulated phylogenies, single cell lineage barcodes, Phylotime reconstructed trees for all experiments..
